# The role of cohesin loading at enhancers in the flux of loop extrusion and long-range transcriptional control

**DOI:** 10.64898/2026.01.20.700462

**Authors:** Erika C. Anderson, Hadi Rahmaninejad, Abrar Aljahani, Emily M. Arnold, Annie S. Adachi, Rini Shah, Karissa L. Hansen, Ivana Cavka, Alistair N. Boettiger, Geoffrey Fudenberg, Elphège P. Nora

## Abstract

Enhancers have been proposed to act as privileged loading sites for cohesin, raising the idea that they actively fold the genome to engage distal target promoters for transcription. Supporting this idea, NIPBL/MAU2, which is required for cohesin loading, binds at enhancers in mouse embryonic stem cells. However, we find that driving cohesin recruitment near an enhancer strongly inhibits transcription from its target distal promoter, indicating that strong focal cohesin loading at enhancers is not compatible with their long-range regulatory functions. Quantitative experiments and biophysical modeling further indicate that cohesin loading at enhancers does not make major contributions to genome-wide cohesin binding and chromosome folding patterns. Instead, cohesin must load throughout the genome to extrude it, regardless of enhancer proximity, with the major determinants of cohesin traffic being extrusion barriers such as transcription and clustered CTCF sites. These findings indicate that enhancer function is largely ancillary to the general mechanisms of chromosome folding, informing further study of the relationship between genome architecture and transcriptional regulation.

## Introduction

Enhancers are genetic elements able to modulate the activity of target promoters over long genomic distances. Such regulatory communication is supported by chromatin loops that bring distal loci into physical proximity within the cell nucleus^1,2^. The cohesin complex plays a key role in this process by dynamically extruding chromatin loops^3,4^. Yet, how the flux of cohesin loop extrusion is controlled to support long-range enhancer-promoter communication remains unclear.

Several studies have proposed that enhancers themselves may serve as loading sites for cohesin^5–12^, thereby directly initiating chromatin loops to contact distal promoters (Fig. 1A). In this view, enhancer function would extend to the reshaping of chromosome folding in ways that support the delivery of their regulatory activity to distal promoters.

**Figure 1:**
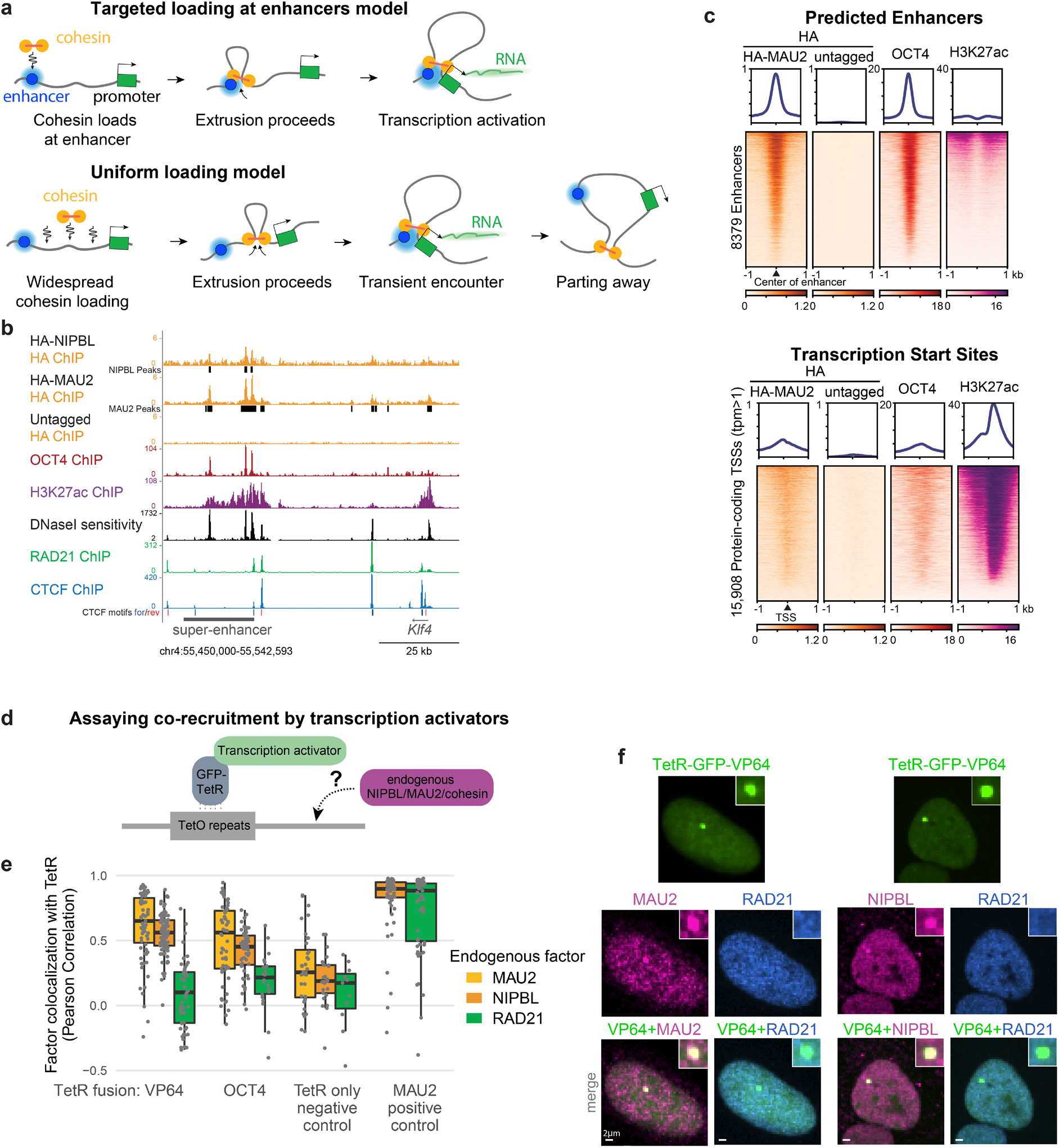
Enhancers bind NIPBL/MAU2 without strongly enriching cohesin. **a**, In the targeted loading model, cohesin is preferentially loaded at enhancers and extrudes toward target promoters to support long-range transcriptional activation. In contrast, in the uniform loading model most cohesin loads away from enhancers and bridges enhancers to their target promoters only transiently. **b**, ChIP-seq in tagged mouse embryonic stem cells (ESCs) shows high NIPBL/MAU2 binding at a distal enhancer, overlapping OCT4 binding sites, but not at a promoter. OCT4 ChIP-seq from ref^51^ and DNAseI sensitivity from ref^115^. **c**, MAU2 ChIP signal is enriched at predicted pluripotency enhancers (from ref^39^) more than that at transcriptionally active TSSs. **d**, Experimental strategy to test co-recruitment of endogenous NIPBL, MAU2, and cohesin by transcriptional activators. TetR-GFP-fused transcription activators were expressed in U2OS cells with a large heterozygous TetO array^40^ and endogenous NIPBL, MAU2, or cohesin were stained by immunofluorescence. **e**, Quantification of colocalization between TetR-GFP and NIPBL, MAU2, or cohesin at the TetO array. Each point denotes the Pearson correlation coefficient between the two proteins in a single cell. TetR-GFP fused to transcriptional activators (VP64 or OCT4) recruits MAU2 and NIPBL but not the cohesin subunit RAD21. TetR alone serves as a negative control, whereas TetR-MAU2 provides a positive control for cohesin recruitment. Boxes indicate interquartile range; center lines denote medians. **f**, Example cells showing that tethering VP64 recruits endogenous MAU2 and NIPBL but no detectable cohesin RAD21.

Support for targeted cohesin loading at enhancers initially came from the observation that, in mammalian genomes, cohesin binds distal regulatory elements along with NIPBL/MAU2^10,13–18^. NIPBL/MAU2 was originally defined as the cohesin loader complex, as it is required to initiate (but not maintain) topological cohesin binding during sister chromatid cohesion^19,20^. In the context of loop extrusion, NIPBL/MAU2 is required for both initiation and extension of extrusion^21–24^. The presence of NIPBL/MAU2 has therefore been interpreted as an indication of cohesin loading and looping from enhancers^10,13,15^. This interpretation was reinforced by the observations that enhancers can locally increase cohesin occupancy^8,11^ and create characteristic chromosome folding signatures such as Hi-C jets/fountains and stripes across organisms^10,25–30^.

However, for loading at enhancers to support long-range looping with distal promoters, cohesin would need to remain stably anchored at the enhancer during extrusion; otherwise cohesin would extrude away from the enhancer before reaching the target promoter (Fig. 1A). Yet, evidence for systematic anchoring of cohesin at enhancers is lacking.

At the same time, alternative models have suggested that cohesin loading is not restricted to discrete NIPBL/MAU2 binding sites (Fig. 1A), but may instead occur in a largely uniform fashion across the genome. Indeed, quantitative simulations of chromosome folding better fit Hi-C and imaging results (including stripes) when they assume uniform rather than targeted cohesin loading^31–33^. Furthermore, many enhancers regulate promoters several hundreds of kilobases away^34–36^, at distances that greatly exceed the average processivity of cohesin molecules that would load at enhancers. Such distances could be bridged by multiple cohesins that load at different loci, but not by exclusive loading at the enhancer. Finally, cohesin loading cannot simply be restricted to topologically associating domains (TADs) containing enhancers, as constitutive heterochromatin is also extruded^37^.

Together, these considerations raise the question to what extent enhancers actively shape chromatin interactions by acting as the loading sites for extrusive cohesin, versus simply operating within the ongoing flux of cohesin extrusion. Despite the relevance to our understanding of the coupling between transcriptional regulation and chromosome folding, discriminating between these possibilities has proven challenging, as it requires interpreting steady-state measurements of cohesin binding and Hi-C within mechanistic models of extrusion dynamics.

Here, we investigate the idea that long-range transcriptional regulation in mammalian cells is facilitated by loading of extrusive cohesin at enhancers through the recruitment of NIPBL/MAU2. We find that, while transcription factors can recruit NIPBL/MAU2 to enhancers, most cohesin molecules load outside enhancers. Artificially recruiting MAU2 to increase cohesin loading and looping near enhancers fails to support distal promoter activation and is instead detrimental to transcription, indicating that highly targeted cohesin loading is incompatible with long-range regulation by enhancers. Consistent with this, genome-wide analyses and quantitative biophysical simulations show that targeted loading at enhancers contributes little, if at all, to the general flux of cohesin extrusion across chromosomes. Instead, cohesin positioning genome-wide is best explained by pervasive loading combined with blocking by transcription and clustered CTCF sites. Together, these results indicate that enhancers do not initiate cohesin loops to support their own action in long-range transcriptional control, and that instead their regulatory opportunities are largely subordinate to the global dynamics of cohesin loop extrusion.

## Results

### Transcription factors recruit NIPBL/MAU2 at enhancers without substantially enriching cohesin

Enhancers have initially been proposed to act as loading sites for extrusive cohesin because they can bind NIPBL/MAU2^10,15^. Given the reported technical difficulties with NIPBL ChIP-seq^31,38^, we first sought to identify high-confidence NIPBL and MAU2 binding sites. We performed spike-in calibrated ChIP-seq with an HA antibody in mouse embryonic stem cells (ESCs) with either endogenous NIPBL or MAU2 tagged with both an HA epitope tag and a dTAG-inducible degron and identified ∼24,000 binding sites (Extended Data Fig. 1A-B). These binding sites were absent in cells where NIPBL or MAU2 was degraded and in cells without an HA tag, ruling out any false positives resulting from antibody binding nonspecifically or to hyper-ChIPable open chromatin (Extended Data Fig. 1B). As expected, HA-NIPBL and HA-MAU2 binding closely matched each other (Extended Data Fig. 1B), and ChIP-seq with an antibody for endogenous NIPBL showed binding at these same sites (Extended Data Fig. 1C). Although these NIPBL and MAU2 ChIP peaks are reproducible, the ChIP signal (with multiple different antibodies) displayed much lower enrichment above background compared to ChIP for other factors (Extended Data Fig. 1G-H). This low enrichment suggests that a substantial amount of the NIPBL/MAU2 molecules bind – and possibly function – outside the peaks.

We analyzed the genomic locations of NIPBL/MAU2 binding, and found that NIPBL/MAU2 was enriched at predicted enhancers (defined as sites bound by the pluripotency transcription factors OCT4, SOX2, and NANOG^39^). In contrast, binding was much less pronounced at active transcription start sites, which show high enrichment for H3K27ac but not pluripotency transcription factors (Fig. 1B-C, Extended Data Fig. 1D-E). Therefore, NIPBL/MAU2 does not simply bind all active chromatin but tracks the binding of pluripotency transcription factors.

To further explore the specificity of NIPBL/MAU2 binding at enhancers, we tested whether transcription factor binding is sufficient to recruit NIPBL/MAU2 and load cohesin (Fig. 1D). We used human U2OS cells with a microscopically visible TetO array spanning ∼4 Mb^40^, a recruitment platform large enough for loaded cohesin to not simply extrude away. Tethering OCT4 and VP64 to the TetO array was sufficient to recruit endogenous NIPBL and MAU2 (Fig. 1E-F, Extended Data Fig. 2) indicating that the ChIP-seq enrichment of NIPBL/MAU2 at enhancers in ESCs is indeed due to recruitment by transcription factors (directly or indirectly via transcriptional cofactors), and that NIPBL/MAU2 recruitment is not limited to pluripotency factors.

Cohesin did not, however, localize to the TetO array upon OCT4 or VP64 recruitment, despite the presence of NIPBL/MAU2. The lack of detectable cohesin upon transcription factor tethering suggests that NIPBL/MAU2 binding to active regulatory elements is not necessarily molecularly coupled to local cohesin recruitment. The low amount of NIPBL/MAU2 recruited by transcription factors is therefore insufficient to cause detectable cohesin accumulation, or NIPBL/MAU2 is not readily competent for loading cohesin when recruited by transcription factors. Recruiting larger amounts of MAU2 by direct fusion to TetR visibly enriched cohesin (Fig. 1E, Extended Data Fig. 2A-B) – demonstrating that upon sufficient loading, cohesin is readily detected by this assay.

### Very strong cohesin loading sites are incompatible with regulation by enhancers

After confirming that NIPBL/MAU2 binds enhancers, we next tested the model that focal cohesin loading at enhancers increases interactions with distal promoters and benefits long-range transcriptional activation. We used the TACL system^41^ to force cohesin loading at the *Car2* locus in ESCs (Fig. 2A), which depends on cohesin for transcriptional activation by distal enhancers^42^. In a cell line where one *Car2* allele encodes a GFP reporter while the other allele encodes RFP (Extended data Fig. 3A), we heterozygously inserted a TetO array either beside the promoter, between the promoter and enhancers, or beside the enhancers, and expressed TetR-MAU2 to target cohesin loading at the TetO array (Fig. 2B). In contrast to the expectation that increasing cohesin loading would bolster activation by the distal enhancers, we observed strong downregulation of *Car2* upon forcing cohesin loading from any of the three TetO locations (Fig. 2B, Extended Data Fig. 3B). Reporter expression was unaffected by TetR alone binding to the TetO array, and expression of the *Car2* reporter on the control allele without a TetO insertion was unchanged (Extended Data Fig. 3C), ruling out possible indirect effects of MAU2 overexpression. Thus, despite the dependence of *Car2* expression on loop extrusion^42^, strong cohesin loading at the enhancer impairs transcriptional activity.

**Figure 2:**
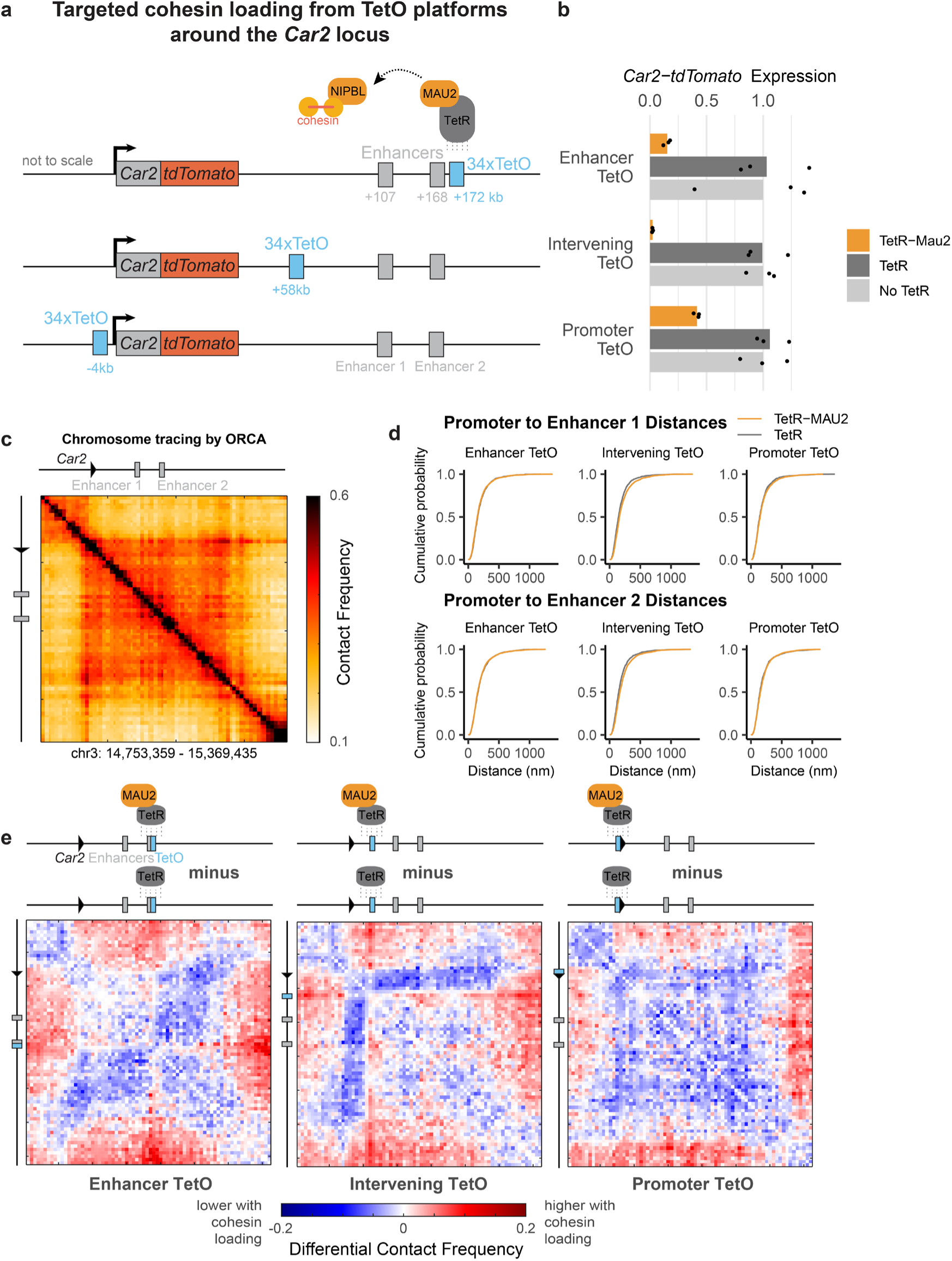
Targeted cohesin recruitment interferes with transcription. **a**, TetR-MAU2 was stably expressed to force cohesin loading at 34x TetO arrays (2 kb) inserted at three positions across the *Car2* locus in ESCs. **b**, *Car2* expression, measured by flow cytometry, was reduced by forcing cohesin loading at all tested positions, whereas TetR binding alone had no detectable effect. Values are normalized to corresponding parental cells. Points show three biological replicates from separate transfections of the respective TetR-expressing plasmid. **c**, DNA FISH chromosome tracing (ORCA) heatmap showing contact frequencies (within ∼200 nm, see methods) at the *Car2* locus (8 kb resolution). **d**, Cumulative plots showing the distribution of 3D distances between the *Car2* promoter and each of its enhancers on the TetO allele with either TetR-MAU2 (forced cohesin loading) or TetR expression (control). **e**, Differential contact maps comparing TetR-MAU2 cells to TetR-only controls (comprising >1300 traces per condition). Targeted cohesin loading does not increase enhancer-promoter interactions but instead creates a boundary, (except at the *Car2* promoter, which is located near an endogenous TAD boundary). The modest reduction in interactions across the TAD arises from global MAU2 overexpression rather than from targeted loading at the TetO site (see allele-specific comparisons within TetR-MAU2 cells in Extended Data Fig. 4B).

To gain insight into the changes in chromatin folding caused by targeted cohesin loading, we used optical reconstruction of chromosome architecture (ORCA, Fig. 2C), which allowed us to specifically probe the *Car2* allele harboring the TetO array (Extended Data Fig. 4A, Methods). Targeting cohesin loading either beside the enhancers or beside the promoter did not increase enhancer-promoter interactions, which remained largely unaffected (Fig. 2D-E, Extended Data Fig. 4C-E). Targeting loading between the enhancers and the promoter separated them by creating a boundary that slightly reduced their interactions (Fig. 2D-E). Therefore, targeted loading-induced transcriptional repression, which occurs from all three configurations, cannot be simply explained by a change in enhancer-promoter interactions. Rather, high levels of extruding cohesin appear to repress transcription by disrupting enhancer or promoter activity, possibly by interfering with assembly or progression of the transcription machinery. Since strong cohesin loading at enhancers is detrimental to transcription from distal promoters, it is unlikely that enhancers generally serve as strong loading sites.

### Polymer simulations show that focal cohesin loading can only moderately impact cohesin traffic

Given our observation that high targeted cohesin loading compromises enhancer function, we sought to establish how more modest loading at NIPBL/MAU2 sites would influence cohesin traffic and chromosome folding. As cohesin at NIPBL/MAU2 sites has been interpreted as a signature of targeted loading^10,43^, we first used biophysical simulations^44^ to explore how much cohesin binding is expected at loading sites across a range of loading rates (Fig. 3A). Using extrusion parameters selected to match measurements in mouse ESCs (Methods), we simulated a uniform background loading rate with elevated loading at a target site and found that bidirectionally extruding cohesin would not accumulate at loading sites, even with up to a 500-fold boost in on-target loading over the background loading rate (Fig. 3B). Instead, cohesin rapidly extrudes away from the loading site.

**Figure 3:**
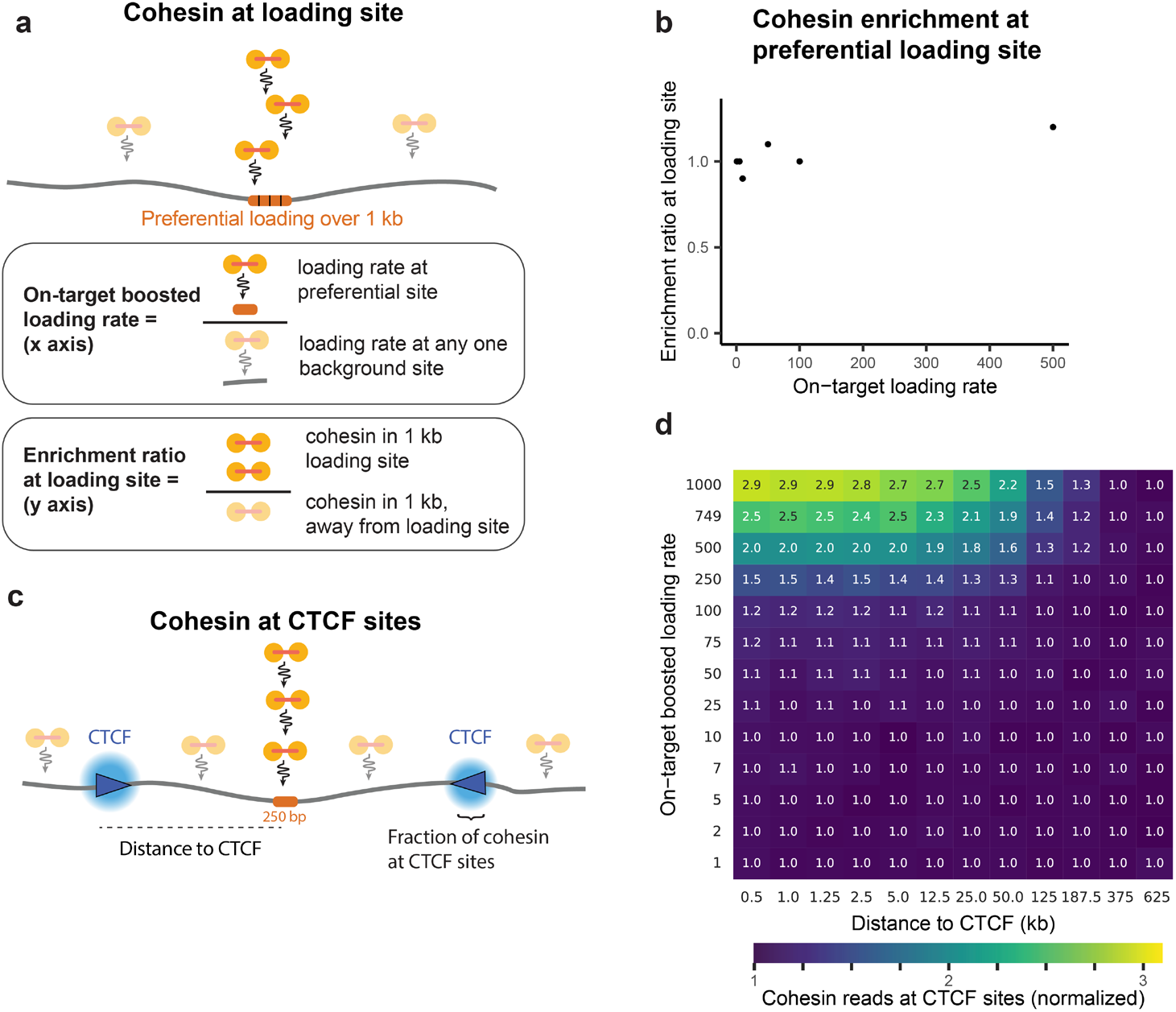
Simulations show minimal effect of targeted loading on cohesin traffic. **a**, and **b,** A 1D lattice simulation indicates that cohesin is not expected to accumulate at loading sites, even at high loading rates. The loading site is modeled as four adjacent monomers spanning 1 kb. The enrichment ratio is defined as the density of cohesin within a 1-kb window centered on the preferential loading site relative to background sites. **c**, Simulations exploring the effect of increasing loading rates at a single site on cohesin accumulation at neighboring CTCF sites (50% occupancy dynamic barrier, see Methods). Here, targeted loading rate at a single 250 bp monomer is increased while maintaining a constant loading rate at all other monomers. **d**, Number of cohesin reads at the CTCF site as a function of the on-target loading boost and the distance between the loading site and CTCF sites. As a reference, values are normalized to the condition with baseline loading rate (1x boost) and a CTCF site positioned 0.5 kb from the loading site.

We next explored to what extent loading sites are expected to increase cohesin accumulation at nearby CTCF sites. To address this, we added to the simulations two dynamic barriers^45^ flanking the loading site at varying distances (Fig. 3C). The simulations revealed that although targeted loading can, in principle, influence cohesin traffic, the impact remains limited. Even for CTCF barriers only 5 kb from the loading site, increasing cohesin accumulation two-fold would require more than a 500-fold boost in on-target loading rate (Fig. 3D). The reason focal loading has limited impact is that the amount of cohesin contributed by targeted loading sites is overshadowed by the cohesins loaded elsewhere along the chromosome. For example, with on-target loading increased 500-fold, a 250 bp loading site would contribute a similar amount of cohesin as background loading from the 125 kb of surrounding chromatin, thereby only increasing cohesin accumulation at the CTCF barrier ∼two-fold. Thus, cohesin traffic to nearby barriers is expected to be dominated by chromosome-wide loading rather than by even strongly privileged loading sites.

Finally, we explored the signatures of targeted cohesin loading on Hi-C patterns. 3D polymer models indicated that high focal loading causes the emergence of fountain patterns in contact maps (Extended Data Fig. 5A-B), as observed in previous simulations ^25,26^. Yet, these patterns do not increase the interactions of the loading site with its surrounding genomic environment (Extended Data Fig. 5A-B, Supplementary video), as cohesin extrudes away from the loading site. We observed that, at high loading rates, wider multi-kilobase loading sites create a boundary with stripes on contact maps (Extended Data Fig. 5B), due to multiple closely loaded cohesins colliding with each other. Although such a boundary at a large loading site could anchor cohesin, our TACL results indicate that such a configuration impairs long-range transcriptional regulation (Fig. 2).

### Most cohesin resides outside CTCF sites

Having defined expectations of how targeted loading can influence cohesin positioning, we next quantified cohesin localization in ESCs to determine the extent to which NIPBL/MAU2 sites at enhancers contribute to initiating loop extrusion. In vertebrates, core cohesin subunits exhibit prominent ChIP peaks at CTCF sites^46–48^, where loop extrusion is blocked. However, cohesin ChIP-seq also exhibits diffuse signal between CTCF peaks, which may reflect true translocating cohesin or non-specific background from the antibody^49^. This ambiguity complicates quantifications, as exemplified by antibodies against distinct cohesin subunits yielding markedly different fractions of reads at CTCF peaks (12% for RAD21 antibody vs. 5% for SMC1A antibody, Fig. 4A-B).

**Figure 4:**
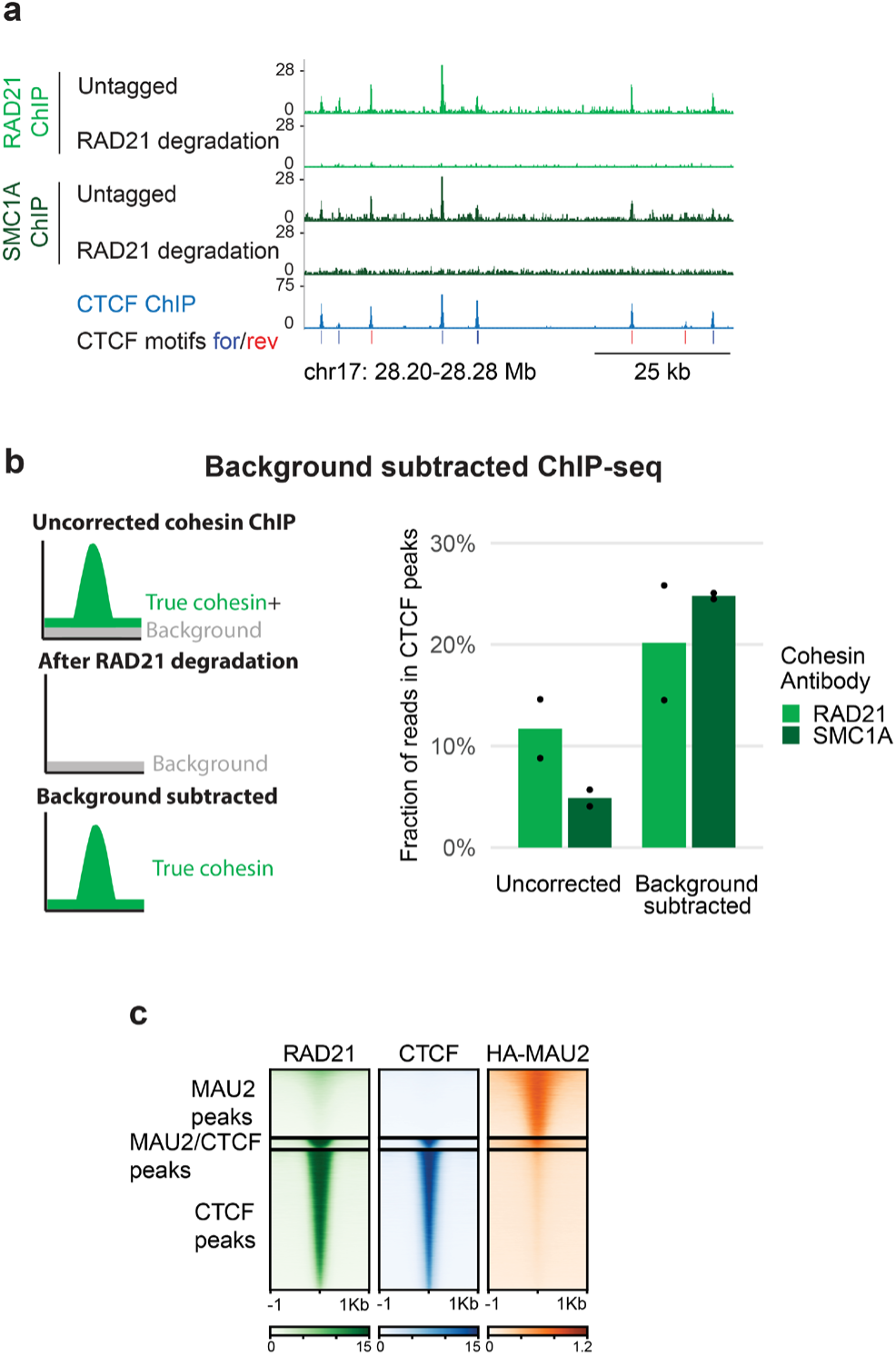
Cohesin binding is largely between CTCF sites. **a**, Spike-in calibrated ChIP-seq for cohesin subunits RAD21 and SMC1A in untagged ESCs or after cohesin degradation in RAD21-AID ESCs treated with auxin (IAA) for 3.5 hours. **b**, Strategy to calculate true cohesin binding by subtracting background antibody signal from the total reads. After background subtraction, CTCF peaks contain ∼25% of cohesin ChIP-seq reads (average of two biological replicates). **c**, ChIP-seq shows little binding of the cohesin subunit RAD21 at NIPBL/MAU2 sites compared to CTCF sites.

To determine how much of the detected ChIP signal is due to nonspecific antibody binding, we performed spike-in calibrated cohesin ChIP after cohesin depletion (Fig. 4B). We depleted RAD21 with an auxin-inducible degron^50–52^ for 3.5 hours (Extended Data Fig. 6A-B) and performed ChIP-seq for RAD21 and SMC1A. This allowed quantifying and subtracting the antibody background from the ChIP-seq signal obtained in cells with normal cohesin levels.

Using this approach, we found that only ∼25% of the cohesin ChIP-seq signal is in the 0.8% of the genome bound by CTCF (Fig. 4B). This background subtraction procedure yielded strong concordance between the RAD21 and SMC1A antibodies, despite their different specificities prior to correction (Fig. 4B). We note that 25% is an upper bound for the fraction of cohesin binding in CTCF peaks, as signal remaining after auxin treatment also includes any undegraded bound cohesin (less than 4%, Extended Data Fig. 6A-B). Thus, the majority of cohesin molecules are positioned between CTCF peaks, where they are likely engaged in extrusion, transiently bound at loading sites, or stalled by collisions or other barriers, such as the transcriptional machinery.

Cohesin enrichment at NIPBL/MAU2 sites was only ∼2-fold over background (Fig. 4C; Extended Data Fig. 6C-D), and most NIPBL/MAU2 peaks did not coincide with RAD21 peaks (Extended Data Fig. 1F). Since our simulations indicated that isolated loading sites are not expected to retain cohesin, we next asked whether NIPBL/MAU2 binding at enhancers nonetheless increases cohesin levels in the surrounding chromatin.

### Proximity to a NIPBL/MAU2 binding site only mildly boosts cohesin levels

Guided by our simulations, we quantified RAD21 ChIP-seq enrichment at CTCF peaks, reasoning that the cohesin level, normalized to CTCF occupancy, reflects the amount of extruding cohesin blocked at each site. While CTCF and RAD21 signals were correlated, the correlation was lower than between CTCF replicates or between RAD21 replicates (Fig. 5A-B, Extended Data Fig. 6E). Accordingly, the RAD21/CTCF ratio varied 15-fold between sites (Fig. 5C), highlighting that CTCF sites with comparable occupancy block markedly different cohesin amounts across the genome (Fig. 5D). This variability is not simply explained by CTCF motif architecture, as cohesin levels were independent of the presence of upstream or downstream auxiliary motifs beyond the core CTCF motif^53^ (Extended Data Fig. 6F).

**Figure 5:**
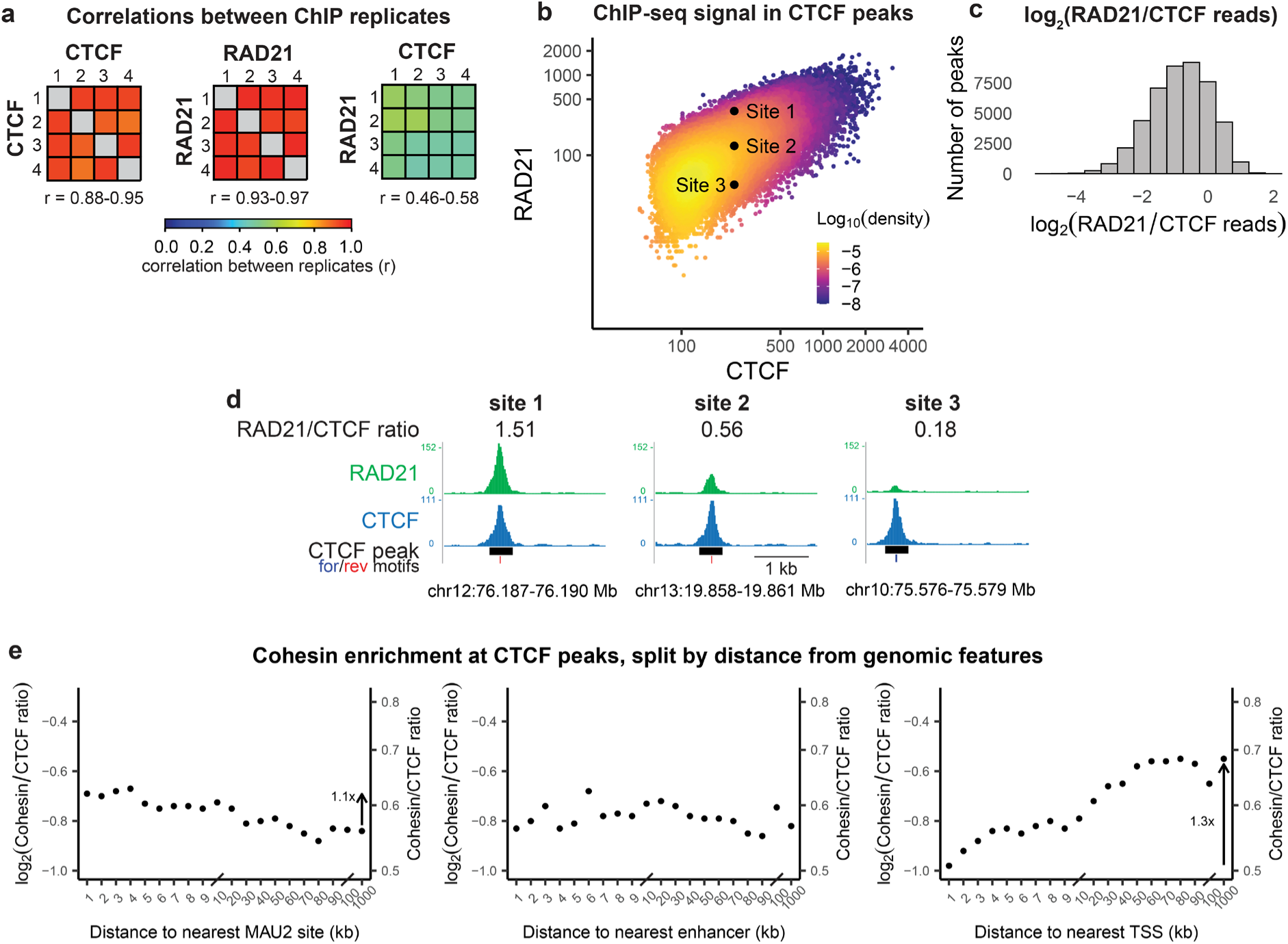
NIPBL/MAU2 binding sites have minimal effects on local cohesin levels. **a**, Correlation matrices (Pearson correlation coefficients) comparing ChIP–seq signal at CTCF sites across biological replicates for CTCF and RAD21, showing that replicates of the same factor correlate more strongly with each other than with the other factor. **b**, Scatter plot of RAD21 vs. CTCF reads in each CTCF peak (from four combined ChIP-seq replicates), highlighting substantial variability in cohesin occupancy at sites with similar CTCF binding. **c**, Distribution of RAD21/CTCF ratios. **d**, Example sites with the same CTCF occupancy but six-fold range in cohesin levels. **e**, Average cohesin/CTCF ratio as a function of the distance from the CTCF peak to the nearest MAU2 site, enhancer^39^, or active TSS, showing minimal dependence on proximity to NIPBL/MAU2 binding and increased cohesin levels at greater distances from active TSSs.

Proximity to NIPBL/MAU2 sites or enhancers did not appreciably influence the RAD21/CTCF ratio: CTCF peaks within 1 kb of a NIPBL/MAU2 site only showed 1.1-fold higher cohesin enrichment compared to sites over 100 kb away from the nearest NIPBL/MAU2 binding site (Fig. 5E). We reached the same conclusion by analyzing ChIP-seq from neural progenitor cells (NPCs, Extended Data Fig. 7A-B), indicating that NIPBL/MAU2 binding sites only have a minimal influence on the amount of cohesin that extrudes to neighboring CTCF sites.

As an orthogonal measure, we inspected whether NIPBL/MAU2 binding sites create chromatin folding signatures expected from targeted cohesin loading, such as Hi-C fountains or stripes^25–27,29,30^ (Extended Data Fig. 5B). Individual strong NIPBL/MAU2 sites did not display such patterns (Extended Data Fig. 8A), although a weak fountain-like signal is present in aggregate heatmaps^25^ (Extended Data Fig. 8B). This observation further supports that cohesin loading at NIPBL/MAU2 sites does not have a major role in shaping chromosome folding patterns in ESCs.

Altogether, our analyses indicate that the cohesin that loads at enhancers is not a major contributor to the flux of loop extrusion. Because other studies reported that enhancer insertions can increase cohesin levels at nearby CTCF sites^8,11^, we considered alternative mechanisms by which enhancers could increase cohesin flux. One possibility is that, instead of acting as direct loading sites, enhancers could increase cohesin recruitment over their broader genomic neighborhood, for example by shifting chromatin domains into an accessible state more permissive to loading. Consistent with this idea, simulations showed that increasing diffuse loading over a 50 kb domain by only 2-fold enriches cohesin at flanking CTCF sites as effectively as increasing targeted loading at a single site 500-fold (Fig. 3D and Extended Data Fig. 5D). Thus, our genomic analyses are consistent with a model where enhancer activation triggers modest, domain-wide modulation of cohesin loading that elevates extrusion flux, without necessarily acting as the direct initiation sites for extrusion.

### Active Transcription and clustered CTCF sites impede cohesin traffic

Together, our experimental observations and simulations demonstrate that targeted loading can only explain a small portion of the 15-fold variation in cohesin enrichment measured at CTCF sites (Fig. 5). We therefore investigated to what extent this heterogeneity in cohesin traffic arises from the location of extrusion barriers, rather than cohesin loading.

Intragenic CTCF sites retained the least cohesin when most highly transcribed (Extended Data Fig. 9A), with the lowest cohesin retained at CTCF motifs oriented in the direction of transcription (Extended Data Fig. 9B). This pattern is consistent with transcribing polymerases as mobile barriers that push cohesin during elongation^9,12,31,54–56^, particularly when cohesin can translocate in the direction of transcription without encountering an opposing CTCF.

Cohesin accumulation at CTCF sites was also distinctly influenced by the density and orientation of CTCF binding. CTCF sites within 1 kb of another site bound ∼1.7-fold less cohesin than isolated sites (>1 Mb apart), in both ESCs (Fig. 6A) and NPCs (Extended Data Fig. 7D). Analysis of tandemly oriented CTCF pairs revealed orientation and spacing-dependent effects, with the most pronounced reduction in cohesin accumulation at sites within 30 kb downstream of a co-oriented site (Fig. 6B and 6C, Extended Data Fig. 7E).

**Figure 6:**
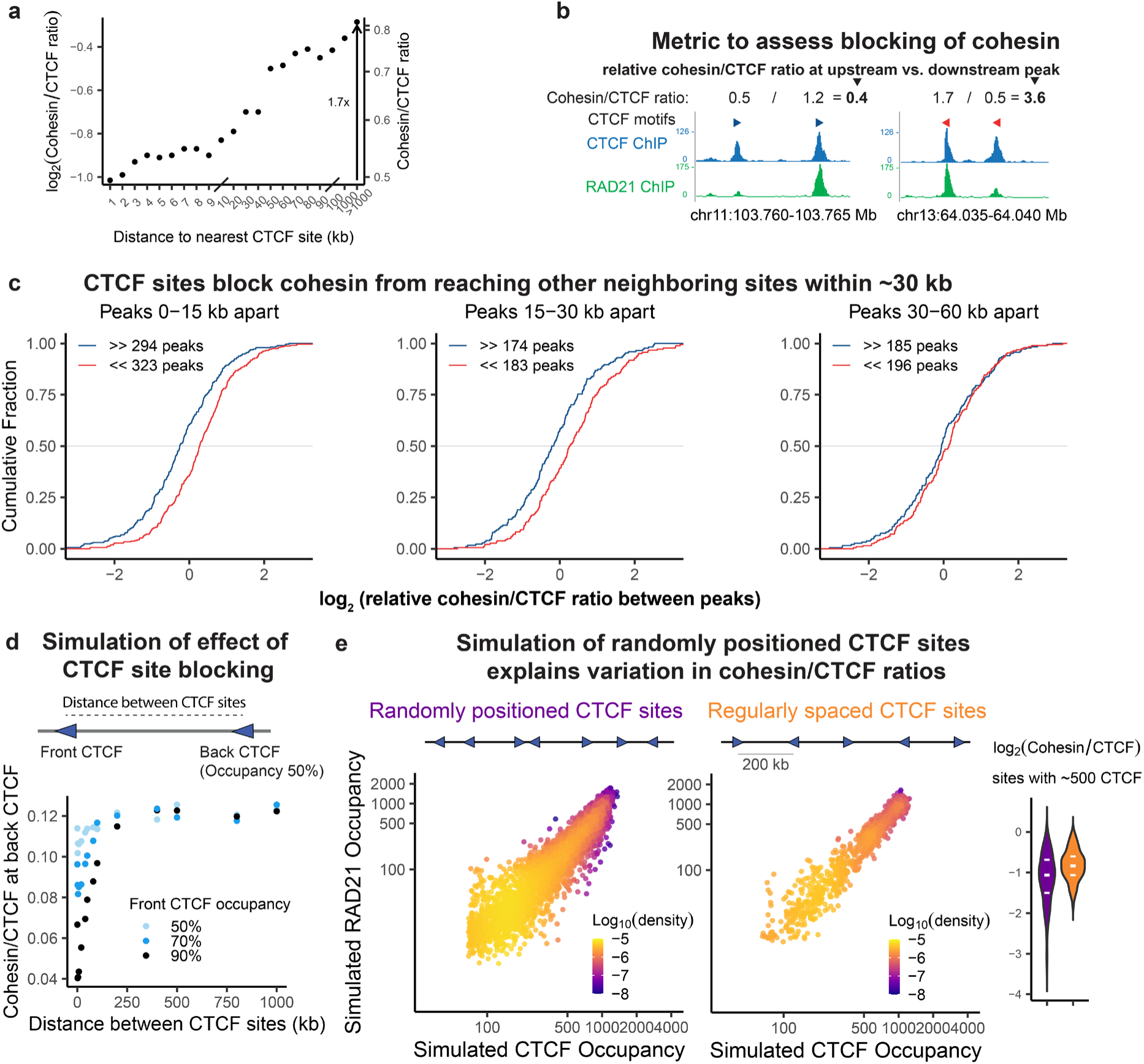
Clustered CTCF sites impede cohesin traffic. **a**, Average cohesin/CTCF ratio at CTCF peaks as a function of distance to the nearest neighboring CTCF site, revealing reduced cohesin occupancy at closely spaced sites. **b**, We identified pairs of CTCF sites with forward motifs and where there is no other CTCF site within the downstream 100 kb. Relative cohesin/CTCF ratios compare upstream and downstream peaks based on motif orientation: a relative ratio less than 1 indicates that the rightmost peak is blocking cohesin from reaching the leftmost peak. For pairs of peaks with reverse motifs, a relative cohesin/CTCF ratio greater than 1 indicates blocking by the leftmost peak. **c**, Cumulative distributions of relative cohesin/CTCF ratios for CTCF site pairs with forward or reverse motifs, stratified by inter-site distance. Interference between neighboring CTCF sites is apparent only for sites within 30 kb of each other. **d**, In a simulation of two CTCF sites with reverse motifs, the cohesin/CTCF ratio at the rightmost site depends on the distance between sites and the occupancy of the left site. **e**, Cohesin vs. CTCF occupancy in a simulation of a 75 Mb chromosome with randomly positioned CTCF sites shows more dispersion than a simulation with CTCF sites regularly positioned every 200 kb. The CTCF occupancy is varied by altering the bound time while keeping the CTCF unbound time constant between sites.

To further explore the mechanistic basis of these genomic correlations, we turned to simulations. We modeled CTCF sites as dynamic barriers where the average occupancy of a site is determined by the duration of CTCF binding (bound time) and the interval between binding events (unbound time)^45^. In simulations of a pair of tandemly oriented CTCF sites, cohesin at the downstream site was reduced up to three-fold when the barriers were immediately adjacent, and increased sharply with increasing spacing (Fig. 6D). High occupancy sites (with longer bound time) blocked most effectively. These results support that barrier interference contributes to the variation of cohesin levels at CTCF sites across the genome.

To explore the interference between CTCF barriers within complex genomic layouts, we simulated a chromosome with many randomly positioned CTCF sites with a range of CTCF occupancies. The resulting cohesin/CTCF ratios showed wide dispersion, with closely spaced CTCF sites retaining the lowest amount of cohesin, recapitulating the patterns seen in ChIP-seq data (Fig. 6E, Extended Data Fig. 10 and 11). Varying either CTCF bound time or unbound time resulted in a wide range of cohesin/CTCF ratios (Extended Data Fig. 10A-B). By contrast, regularly spaced CTCF arrays produced a narrower range of cohesin/CTCF ratios, even when using CTCF sites of variable occupancies (Fig. 6E, Extended Data Fig. 10). These simulations highlight how configurations with clustered barriers can substantially suppress cohesin traffic beyond the effect of any individual site.

Finally, we asked whether such sequence-encoded interference is captured by predictive machine learning models that are mechanism-agnostic. Using Enformer (Avsec et al., 2021), we found that introducing CTCF sequences with high cohesin binding into a recipient locus with flanking CTCF sites lowered predicted cohesin levels, mirroring experimental patterns (Extended Data Fig. 12). Removing the flanking CTCF motifs abolished this effect, illustrating that even sequence-based models implicitly capture interference between neighboring CTCF sites.

Altogether, these findings reveal that genome-wide cohesin positioning is shaped more by interference between extrusion barriers – arising from transcription and clustered CTCF sites – rather than by targeted loading, providing a simple explanation for the wide dispersion of cohesin levels at CTCF sites across the genome.

## Discussion

Together, our findings argue against a model in which targeted cohesin loading at enhancers increases their functional interactions with distal promoters to boost transcription. Indeed, we observed that, at a locus that relies on loop extrusion for enhancer-promoter communication, increasing cohesin loading at the enhancer represses transcription from the target promoter 170 kb away. Recent studies have also reported that targeted cohesin loading causes transcriptional repression of neighboring promoters^41^, not activation – even at physiological loading rates from endogenous enhancers^30^.

Our analyses highlight that, as enhancers only constitute a small fraction of the genome, their contribution to overall cohesin on chromosomes is overshadowed by baseline loading across the rest of the genome, even with high loading rate at enhancers (Extended Data Fig. 13). Yet, our results do not rule out that, in specialized genomic or cellular contexts, focal loading at some enhancers could modulate interactions with neighboring promoters in a functionally relevant way. Indeed, small changes in enhancer-promoter contact frequency can meaningfully affect transcription – especially when the baseline contact frequency is low^57^. Focal loading at the enhancer could increase interactions with a promoter if cohesin remained anchored at the enhancer due to an additional factor such as CTCF binding^8,10,58^, possibly RNA polymerase^12,31^, or, speculatively, factors that may inhibit direction switching of extrusion^59^. However, most ESC enhancers do not display chromosome interaction patterns (fountains) that would be indicative of elevated interactions. Our observations indicate that long-range transcriptional regulation largely operates without relying on cohesin loading and loop initiation at enhancers, even at cohesin-dependent loci.

Several studies have demonstrated that enhancers can influence the local cohesin flux and genome folding, raising the question of how such effects can be reconciled with their minimal contribution to overall cohesin loading. Our observations are consistent with a model where enhancers modulate cohesin levels indirectly, by increasing loading across an entire domain rather than acting as discrete initiation sites for extrusion. Consistent with this view, our simulations show that modest increases in domain-wide loading produce substantially larger effects on cohesin accumulation at CTCF sites than even stronger focal loading. These simulations of a broad, but not focal, boost in cohesin loading can quantitatively account for the observation that inserting an enhancer in a silent domain leads to a three-fold increase in cohesin at a CTCF site over 100 kb away^8^. Furthermore, studies in different cell types have found that inserting an enhancer can create a new self-interacting domain and increase cohesin binding at sites across the domain^8,60^. Additionally, regions with enhancers or other markers of active chromatin display higher domain-wide interactions^26,61^, and regain domain-wide self-interactions faster after transient cohesin depletion^62^. Studies in yeast demonstrated that chromatin remodelers can recruit NIPBL/MAU2^63^, suggesting that more accessible chromatin domains may experience more pervasive cohesin loading.

The idea of mammalian enhancers facilitating domain-wide rather than focal cohesin loading remains compatible with observed Hi-C fountains. Fountains in differentiated mammalian cells^26^ emerge from active regions hundreds of kb in width that are surrounded by heterochromatin. Such fountains are not evident at ESC enhancers in baseline conditions^25^ (Extended Data Fig. 8) - perhaps due to the generally open chromatin of ESCs^64^, which could result in more uniform cohesin loading. However, the fountains that appear in ESCs after depletion of CTCF and the cohesin unloader WAPL are similarly located at multi-kilobase active domains flanked by heterochromatin^29^. Altogether these considerations are consistent with active enhancers inducing recruitment of cohesin across the width of the active domain, rather than acting as the direct initiation sites of extrusion.

Our analyses illustrate how even modest increases in pervasive, domain-wide cohesin loading can substantially elevate the local flux of loop extrusion. By increasing domain-wide cohesin loading, enhancers could increase chances of transient but functionally productive encounters with distal promoters^65^, without the need to directly initiate cohesin loading at levels that risk interfering with transcription (Fig. 1A).

In this context, the functional relevance of NIPBL enrichment at regulatory elements remains to be clarified. NIPBL binding at enhancers has been consistently reported across studies in various cell types^15,66–68^, while NIPBL binding at promoters was only reported by some^15,17,38,54,66,67^ but not all^31^ studies. Our observation that NIPBL binding is enriched at enhancers over promoters in ESCs is explained because most pluripotency transcription factor binding sites are distal to promoters^69^, while in other cell types, NIPBL binds with transcription factors that are found at promoters^38^. NIPBL recruitment likely occurs through direct interactions with cell-type specific transcription factors^70^ or via intermediate co-factors present across cell types, such as Mediator^15,71^, BRD4^72^, or the transcription complex itself^73^. Whether NIPBL has cohesin-independent functions at regulatory elements and whether NIPBL/MAU2 ChIP-seq peaks depend on cohesin loop extrusion requires further clarification, as studies have reported conflicting results^31,41^.

Regarding the presence of cohesin at enhancers, many studies, especially in differentiated cells, reported low but detectable cohesin enrichment at enhancers^5,6,10,15–17,66,68,74–76^. Given our results, extrusion blocking emerges as a more plausible explanation than targeted loading for cohesin enrichment at such lineage-specific transcription factor binding sites^77^. Supporting a blocking-based mechanism, deletion of transcription factor binding sites within an embryonic stem cell enhancer increases cohesin traffic toward neighboring CTCF sites^78^, and polymer simulations better reproduce chromosome folding pattern by implementing blocking rather than simply loading at enhancers alone^12,77^.

Several non-exclusive mechanisms can be envisioned for how enhancers may block cohesin (independently of CTCF). First, the transcription machinery itself can act as a barrier to cohesin extrusion^9,12,31,56^. Second, extrusion may halt upon direct or indirect interaction between cohesin and factors or structures present at regulatory elements. For example blocking at TSSs could be mediated by the chromatin reader protein Phf2, which translocates with cohesin and may arrest extrusion upon binding H3K4me3^79^. Finally, blocking may arise from indirect physical constraints. For example, transcriptional condensates have been reported to exert forces on DNA *in vitro* of up to 7pN^80^, far exceeding the force sufficient to stall extrusion (∼0.1-0.4 pN, ^81^), providing a potential explanation for the low-level and diffuse cohesin enrichment detected by ChIP-seq at enhancers, without the need to invoke direct TF-cohesin interactions^18^. The mechanisms of cohesin blocking at enhancers and the relevance in long-range regulation remain to be investigated.

Our quantitative analyses of cohesin ChIP-seq reveal that most molecules (>75%) are found outside CTCF sites. This result is consistent with orthogonal imaging measurements indicating that TADs are rarely fully extruded even at steady-state^82–85^. By extracting interpretable signatures of extrusion dynamics from snapshot cohesin ChIP-seq, our work explains why cohesin occupancy is a better predictor of 3D genome folding patterns than CTCF occupancy alone^86^: the impact of a CTCF site on cohesin traffic depends on the amount of the extrusion flux it can intercept, which itself depends on the density, orientation, and binding kinetics of nearby CTCF sites. These considerations are directly relevant to the interpretation of CTCF site disruptions (with mutations of isolated CTCF sites producing larger chromosome folding defects^43,87–89)^ and developmental loci where tuning cohesin trajectories implements cellular decisions^33,90,91^.

Finally, our work does not exclude functional roles for targeted cohesin loading, especially in genomic processes beyond enhancer regulation. For example, targeted increases in cohesin loading have been proposed to guide DNA double-strand break repair by non-homologous end joining^92^, a process accompanied by local transcriptional suppression^93^. We speculate that DNA break-induced targeted cohesin loading may contribute to the local suppression of transcription. In *S. cerevisiae,* targeted loading of condensin, another loop extruder, guides recombination events underlying mating type switching^94^.

Taken together, our work reframes the relationship between transcriptional regulatory elements and cohesin biology, illuminating how genomic context shapes the local flux of loop extrusion within which enhancers operate.

## Methods

### Plasmid Construction

Plasmids were assembled using Gibson assembly (NEB HiFi DNA Assembly Master Mix E2621) or restriction-ligation (NEB Quick Ligase M2200). A list of all plasmids used in this study is available in Supplementary Table 1 and 2.

### Cell culture

Parental wild-type E14Tg2a (karyotype 19, XY, 129/Ola isogenic background) *Mus musculus* embryonic stem cells and subclones were cultured in 2iSL medium [DMEM + GlutaMAX + sodium pyruvate (ThermoFisher 10569044) supplemented with 15% fetal bovine serum (Gibco SKU A5256701), 550 µM 2-mercaptoethanol (ThermoFisher 21985-023), 1X nonessential amino acids (ThermoFisher 11140-050), 10^4^ U/mL of Leukemia Inhibitory Factor (Millipore ESG1107), 1 µM PD0325901 (Apex Bio/Fischer A3013-25), and 3 µM CHIR99021 (Apex Bio/Fischer A3011-100)]. Cells were maintained at a density of 0.2-1.5 x 10^5^ cells/cm^2^ by passaging using TrypLE (ThermoFisher 12605010) every 1-2 days on 0.1% gelatin-coated dishes (Sigma-Aldrich G1890-100G in 1XPBS) at 37°C and 5% CO_2_. The medium was changed daily when cells were not passaged. Cells were checked for mycoplasma infection every 6 months and tested negative.

To establish neural progenitors (NPCs), 0.2 million WT E14 or NIPBL-FKBP ESCs were seeded in a gelatinized T25 flask in 2iSL medium. The following day, cells were rinsed twice in 1X PBS and switched to N2B27 medium [50% DMEM/F12 medium (Gibco 31330-038), 50% Neurobasal medium (Gibco 21103-049), 1X GlutaMAX (Gibco 35050061), 0.5X B27 (Gibco 17504-044), 0.5X N2 (Millipore SCM012), and 0.1 mM 2-mercaptoethanol], and media was changed daily. After 7 days, cells were detached using TrypLE, seeded in nongelatinized bacterial dishes for suspension culture at 3 million cells per 75 cm^2^, and cultured in N2B27 containing 10 ng/mL EGF and FGF (Peprotech 315-09 and 100-18B). After 3 days, floating aggregates were seeded on gelatinized dishes. After 2-4 days, poorly attached aggregates were dislodged and discarded, and remaining adherent NPC cells were dissociated using Accutase (Innovative Cell Technologies AT104), passaged twice on gelatinized dishes in N2B27 + EGF/FGF and cryopreserved after expansion in 10% DMSO in N2B27 + EGF/FGF. Cells were thawed and cultured on gelatinized dishes in N2B27 + EGF/FGF and passaged using Accutase for 2-3 minutes at room temperature.

FKBP^12F36V^ depletion was triggered by adding PROTAC dTAG-13 (dTAG) (Sigma-Aldrich SML2601) to a final concentration of 500 nM. AID depletion was triggered by adding indole-3-acetic acid sodium salt (IAA, auxin analog) (Tocris Bioscience 7932) to a final concentration of 500 mM.

HEK cells were grown in DMEM with 10% FBS and split when 70% confluent using TrypLE. The medium was changed every 2-3 days.

U2OS cells were grown in McCoys 5A media (Thermo Fisher 16600082) supplemented with 10% FBS at 37°C and 5% CO2. Cells were passaged every 2-3 days when 70-80% confluent using TrypLE.

### ESC Genome engineering

For transfection, plasmids were prepared using the Nucleobond Midi kit (Macherey Nagel 740410.5). Transfections were done using the Neon system (Thermo Fisher) with a 100 µL tip and 1 million cells at 1400 V, 10 ms, 3 pulses. To create knock-in cells, ESCs were co-transfected with 5 µg of vector expressing Cas9 and sgRNA and 20 µg of targeting construct. Homology arms of targeting vectors were ∼1 kb. For deletions, 2.5 µg each of two sgRNAs targeting the two ends of the deletion were used.

After electroporation, cells were seeded in a 9 cm^2^ well and left to recover for 48 hours. Cells were then plated at limiting dilution and grown for 8-11 days, changing medium every other day, until single colonies could be picked. Individual colonies were genotyped by PCR and validated with Sanger sequencing.

When the targeting construct contained a resistance cassette, sgRNAs were cloned into pX330, for co-expression of the sgRNA and Cas9. Cells were grown with antibiotic in the medium from the time of serial dilution until the colonies were picked (Puromycin: 1 µg/mL; Hygromycin: 200 µg/mL).

When the targeting construct did not contain a resistance cassette, we used sgRNAs cloned into a Cas9-2A-puro vector (pEN243, identical to pX459 Addgene 62988). Cells were treated with puromycin from ∼24-72 hours after transfection to kill untransfected cells.

For mESC experiments using lipofectamine transfections, 360 ng total of plasmid was mixed with 150 ul OptiMEM, and separately 1.8 ul lipofectamine 2000 was mixed with 150 ul OptiMEM. Both mixtures were incubated in the dark for 5 minutes, then combined and incubated for an additional 60 minutes in the dark at room temperature. 300 ul of the mixture was then added to 0.3 million ESCs seeded in a 9 cm^2^ well. A list of all cell lines used in this study is available in Supplementary Table 3.

### HEK cell genome editing

We created a HEK cell line in which the C-terminal of CTCF is tagged with a series of epitopes (HA-FLAG-TY1-eGFP-T2A-PuroR). For genome editing, 1 million cells were transfected with 20 ug of the targeting construct and 5 ug of the Cas9 sgRNA plasmid. We used the Neon system with a 100 µL tip at 1100 V, 20 ms, 2 pulses. After transfection, the cells recovered for four days, with 1 µg/mL puromycin added starting two days after transfection. Four days after transfection, the cells were split, diluted, and dispensed into 96-well plates at a concentration such that there was an average of 0.5 cells per well. After 10 days of growth, single colonies were genotyped by PCR. Of 27 colonies, 10 appeared homozygous for the insertion, and the remainder heterozygous for the insertion. The insertion was verified by sequencing, and GFP expression was confirmed by flow cytometry.

### Western blotting

We used Western blotting to quantify RAD21 and MAU2 depletion. Approximately 10 million ESCs were dissociated using TrypLE, resuspended in culture medium, and pelleted at 100g for 5 minutes at room temperature. Cell pellets were washed in PBS, pelleted again, snap frozen on dry ice after aspirating the PBS, and stored at −70°C. Cell pellets were resuspended in 10 mM HEPES (pH 7.9), 2.5 mM MgCl2, 0.25 M sucrose, 0.1% NP40, 1 mM DTT, and 1X HALT protease inhibitors (ThermoFisher 78437) and kept on ice for 10 minutes to allow swelling. After centrifugation at 500g at 4°C for 5 minutes, the supernatant was discarded and the pellets were resuspended on ice in 25 mM HEPES (pH 7.9), 1.5 mM MgCl_2_, 700 mM NaCl, 0.5 mM DTT, 0.1 mM EDTA, 20% glycerol, 1 mM DTT, and 250 U benzonase (Sigma-Aldrich E1014), and incubated on ice for 10 minutes. The nuclear lysates were centrifuged at 18,000g at 4°C for 10 minutes, to eliminate the cellular debris. Protein concentration from the supernatants was measured using Pierce BCA Kit (ThermoFisher 23225).

40 ug of protein extract was combined with 1x Laemmli buffer (Biorad 1610747), heated at 95°C for 10 minutes, and subjected to electrophoresis on a 4-15% polyacrylamide gel (Biorad 17000927). The separated proteins were transferred onto a polyvinylidene difluoride (PVDF) membrane (Millipore IPFL00010) using phosphate-based transfer buffer (10 mM sodium phosphate monobasic, 10 mM sodium phosphate dibasic) at 4°C, 600 mA, for 1.5 hours. After the transfer, the membrane was cut so the lower molecular weight section could be blotted with TBP antibody for the loading control. Membranes were blocked in 5% blocking buffer (5% skim milk in TBS-T (20 mM Tris-Cl buffer-pH 7.4, 500 mM NaCl and 0.1% Tween 20)) and incubated overnight at 4°C with the primary antibodies (RAD21 Millipore 05-908 1:1000, RAD21 abcam ab154769 1:1000, MAU2 abcam ab183033 1:1000, or TBP Cell Signaling Technology 8515S 1:2000) prepared in 5% blocking buffer. The next day, the membranes were washed three times with TBS-T buffer and incubated with mouse or rabbit secondary antibodies conjugated with horseradish peroxidase diluted 1:2000 (Biotium 20400, Biotium 20403) for 1 hr at room temperature. The membranes were again washed three times with TBS-T buffer. The blots were developed using Pierce ECL western blotting substrate (ThermoScientific 32209) and detected using Chemidoc XRS+ imager (Biorad).

Three biological replicates of untagged (E14) and RAD21-GFP-AID cells with and without auxin treatment were collected on separate days. Intensity of the bands was quantified using ImageJ. For Extended Data Fig. 6A, RAD21 intensities were normalized to TBP and then to the value in untagged cells.

### Flow cytometry

ESCs were dissociated with TrypLE, resuspended in culture medium, pelleted, and resuspended in 10% FBS in PBS before live cell flow cytometry on an Attune NxT instrument (Thermo Fisher). Analyses were performed using the FlowJo software. Briefly, live cells were gated using FSC-A x SSC-A and singlets were then gated using FSC-H x FSC-A before extracting median values for approximately 50,000 final gated cells. The value of autofluorescence in untagged E14 cells was subtracted.

For quantification of RAD21-GFP, values were normalized by dividing by the average expression in untreated cells, and replicates were obtained by collecting cells on different days.

### Calibrated ChIP-seq

For all NIPBL and MAU2 ChIP experiments, cells were double crosslinked in disuccinimidyl glutarate (DSG) followed by formaldehyde. First, cells were dissociated using TrypLE (ESCs) or Accutase (NPCs) and resuspended in culture medium. 15 million cells were pelleted at 100*g* for 5 minutes at room temperature. Supernatant was discarded and cell pellets were washed once in PBS. Cells were resuspended in 10 mL PBS, and DSG (Fisher Scientific AAH58208MC) dissolved in DMSO was added to a final concentration of 2 mM. Cells were fixed at room temperature for 45 minutes with rotation. Cells were then pelleted at 200*g* for 5 minutes and washed three times with PBS (each time pelleting at 200*g* for 5 minutes). Cells were then resuspended in 5 mL of 10% FBS prepared in PBS. To this, 5 mL of freshly prepared 2% formaldehyde solution was added to reach a final formaldehyde concentration of 1%. 2% formaldehyde was diluted from 16% methanol-free formaldehyde (Fisher Scientific PI28908) in 1X PBS with 10% FBS. Resuspended cells were fixed in formaldehyde for 10 minutes at room temperature with rotation. To quench, 0.525 mL of 2.5 M glycine (Millipore Sigma GX0205-1) solution in PBS was added (final concentration 0.125 M glycine), and cells were incubated for 5 minutes at room temperature followed by 15 minutes on ice. Cells were pelleted by centrifuging at 200*g* for 5 minutes at 4°C. Cell pellets were resuspended in 1 mL of ice-cold 0.125M glycine in PBS, transferred to 1.5 ml tubes, and centrifuged at 300*g* for 5 minutes at 4° C. Supernatant was discarded and pellets were snap frozen on dry ice and stored at -70°C.

For CTCF and RAD21 ChIP, cells were dissociated, pelleted, and fixed in 1% formaldehyde for 10 minutes with the subsequent quenching and washes as above.

For ChIP, fixed cells were thawed on ice, resuspended in 1 mL of ice-cold lysis buffer [10 mM Tris-HCl pH 7.5, 10 mM NaCl, 0.2% NP-40 (IGEPAL), 0.2% TritonX-100, 1 mM EDTA, 0.5 mM EGTA, supplemented with fresh 1x protease inhibitor (Thermo Scientific PI78437)] and incubated on ice for 10 minutes. ∼9.75 x 10^6^ mouse nuclei were combined with ∼0.25 x 10^6^ fixed HEK nuclei as spike-in. For ChIP experiments with HA antibody, HEK cells with an HA epitope (cell line EA46.3) were used for the spike-in. Nuclei were pelleted at 400*g* for 5 minutes at 4°C, the supernatant was discarded, and nuclear pellets were resuspended in 1 mL of shearing buffer (0.5% SDS, 10 mM Tris-HCl pH 7.5, supplemented with fresh protease inhibitor). Chromatin was sheared using an S220 Covaris with settings PIP 105, duty factor 5, cycles per burst 200, total time 23 minutes for double-crosslinked cells or 15 minutes for formaldehyde-crosslinked cells. After shearing, samples were spun at 13,000 rpm for 5 minutes at 4° C. 2.5% of the sample was removed to use as input, and the remainder of the supernatant was used for ChIP. The ChIP reaction was assembled as follows: 900 ul chromatin, 442 μL 5X ChIP incubation buffer (5% TritonX-100, 0.75 M NaCl, 5 mM EDTA, 2.5 mM EGTA, 50 mM Tris-HCl pH7.5), 40 uL Dynabeads (pre-rinsed in ChIP incubation buffer), 10 ug antibody, 22.5 uL protease inhibitor, and 835.5 uL water. Tubes were incubated overnight at 4°C on a rocker.

The following antibodies were used for ChIP: HA (Thermo Fischer 26183), RAD21 (Abcam ab992), RAD21 only for experiments in RAD21-AID cell lines (abcam ab154769), CTCF (Active Motif 61311), NIPBL (Bethyl A301-779A), H3K27ac (abcam ab4729).

Following overnight incubation, beads were collected using a magnetic rack and washed sequentially with different chilled wash buffers for 5 minutes each on a rocker at 4°C. Beads were washed twice with 500 μL of wash buffer 1 (0.1% SDS, 0.1% Sodium deoxycholate, 1% TritonX-100, 0.15 M NaCl, 1 mM EDTA, 0.5 mM EGTA, 10 mM Tris pH 8), once with 500 μL of wash buffer 2 (0.1% SDS, 0.1% Sodium deoxycholate, 1% TritonX-100, 0.5 M NaCl, 1 mM EDTA, 10 mM Tris pH 8), once with 500ul of wash buffer 3 [0.25 M LiCl, 0.5% Sodium deoxycholate, 0.5% NP-40 (IGEPAL), 1 mM EDTA, 0.5 mM EGTA, 10 mM Tris pH 8], and twice with 500 μL of wash buffer 4 (1 mM EDTA, 0.5 mM EGTA, 10 mM Tris pH 8).

To elute the DNA, beads were resuspended in 200 μL of elution buffer (1% SDS, 0.1M NaHCO_3_) and incubated on a rocker for 20 minutes at room temperature. Beads were then separated using a magnetic rack, and the eluate was transferred to a fresh tube while beads were discarded. Input samples were thawed and elution buffer was added to bring the volume to 200 μL. 2 μL of 1 mg/ml RNaseA (Thermo Scientific FEREN0531) was added before incubation at 37°C for 30 minutes. To reverse crosslinks, 8 μL of 5 M NaCL and 5 μL of 10 mg/ml Proteinase K were added, and samples were incubated overnight at 65°C. DNA was eluted using a MinElute kit (Qiagen 28604).

Libraries were prepared using NEBNext Ultra II DNA library prep kit according to the manufacturer’s protocol (E7645S). The entire ChIP sample and 50 ng of the input samples were used for library prep. Libraries were amplified for 16 cycles, followed by 0.9X SPRI bead purification (Bulldog Bio CNGS005). The eluate was run on a 2% E-gel (Invitrogen G401002) and fragments between 200-600 bp were excised and eluted using a MinElute gel extraction kit (Qiagen 28604). Quality and quantity of libraries were assessed using Qubit and Bioanalyzer High Sensitivity DNA kit (Agilent 5067-4626). Libraries were sequenced on a NextSeq 2000 using 65 bp single-end reads. Some libraries were sequenced in a second lane for additional reads. Each replicate is a population of cells fixed on a separate day.

### ChIP-seq mapping

Reads from fastq files were aligned to a catenated mm10 and hg38 reference genome using Bowtie2. Reads with a mapq score ≥ 30 were retained and duplicates were removed, using samtools version 1.21 ref^95^. ChIP bigwigs were generated with the bamCoverage function from deepTools^96^, with scaling factors calculated using the formula (human reads in input) / (mouse reads in input) / (human reads in ChIP) x 15,000,000 ref^97^. The multiplier of 15,000,000 was chosen arbitrarily to set the final scaling factors close to 1. Input bigwigs were normalized using RPKM. Bigwigs were visualized using the UCSC genome browser^98^.

To create bigwig files with multiple replicates combined, we took equal numbers of reads from each replicate (samtools view with subsample option), combined them, and generated a bigwig normalized by RPKM.

### Creating consensus ChIP-seq peak lists

To create lists of ChIP-seq peaks found in multiple replicates, we first used macs2 version 2.2.9.1^99^ to call peaks on each individual ChIP replicate. For NIPBL and MAU2 peaks, we used HA ChIP in untagged cells as the control. For CTCF and H3K27ac peaks, we used the corresponding input sample as the control. We used bedtools multiinter to identify regions that were called as peaks in the majority of replicates. To define the peak widths, we again called peaks using macs2, this time including all of the replicates together. Using bedtools intersect -u, we selected the peaks from the list called using combined replicates that overlap with the regions called in the majority of replicates. Finally, we removed peaks in ENCODE blacklisted regions^100^ using bedtools intersect -v.

All analyses of NIPBL/MAU2 peaks use the consensus list of HA-MAU2 peaks, as the HA-MAU2 ChIP was more sensitive compared to the HA-NIPBL ChIP.

### Comparing ChIP-seq binding

We generated tornado plots and plots of average binding at ChIP-seq peaks using computeMatrix and plotHeatmap functions from deepTools. OCT4 ChIP-seq is from ref^51^, file name GSM3992929_33_WaplC6-0h_antiOct4.bw, and H3K4me3 ChIP-seq is from ENCODE mouse ES-E14 dataset ENCSR000ADL, file ENCFF001MXU, both mapped to mm10 as above. We used the H3K4m3 peak list ENCFF012AZO. We generated the upset plot (Extended Data Fig. 1E) using Intervene^101^.

### TetO array imaging

We recruited transcription factors to a TetO array in U2OS 2-6-3 cells^40^. We avoided using the LacO repeats that are also present in the array as we observed that LacR expression alone led to cohesin recruitment, possibly because of chromosomal instability triggered by the tighter LacO-LacR binding^102^. In eight-well ibidi slides (Fisher 50305795), we seeded 20,000 cells per well and transfected with a plasmid expressing either TetR-VP64 (pEM037), TetR-OCT4 (pEM040), TetR (pEM041), or TetR-MAU2 (pAA014) using lipofectamine 3000 (Thermo Fisher L3000008) according to the manufacturer’s instructions.

For immunofluorescence, we rinsed cells once with 250 uL of PBS and then fixed in 3% formaldehyde (Electron Microscopy Sciences 15686) diluted in PBS for 10 minutes at room temperature. Cells were rinsed three times with PBS and permeabilized with fresh 0.5% Triton X-100 in PBS for 5 minutes. After rinsing three times with PBS, cells were incubated with 1% BSA in PBS for 10 minutes. Next, cells were incubated in a 150 uL volume of primary antibody diluted 1:300 in 1% BSA in PBS for 45 minutes (antibodies: NIPBL Bethyl A301-779A, MAU2 abcam ab183033, RAD21 Millipore 05-908). Cells were washed five times with 250 uL PBS for 5 minutes each. Next, cells were incubated with secondary antibody diluted 1:1000 in 1% BSA in PBS for 45 minutes (goat anti-rabbit Invitrogen A-11012, goat anti-mouse Invitrogen A-11005) and again washed three times. Finally, cells were incubated with 1 ug/mL DAPI in PBS for 5 minutes and rinsed twice in PBS. All steps were performed at room temperature.

Images were acquired using a Zeiss spinning disk confocal microscope with 60x objective or Nikon spinning disk confocal microscope with a 60x objective with XY pixel size of 183 x 183 nm and 300 nm steps in Z. Images were analyzed in ImageJ with the JACoP plugin to calculate the Pearson correlation between red and green channels within a 12 × 12 × 8 X × Y × Z box manually placed on each GFP-positive TetO array. To calculate the intensity of signal at the array compared to background, for each cell we extracted the total signal within a box centered on the array of size 6x6 XY pixels and 4 z slices. We then chose a box outside the array with a representative level of cohesin of size 12x12 XY pixels and the same 4 z slices as the first box. To account for the different box sizes, we computed enrichment at the array as the (array intensity * 4) / (background intensity).

### TACL

We heterozygously inserted a 34x TetO array at different locations in a cell line in which the two alleles of *Car2* are tagged with tdTomato and mClover reporters. To facilitate distinguishing between the two alleles by ORCA, we first created a 10 kb deletion of a neutral sequence near *Car2* enhancer 2 (chr3:15062000-15072751).

We then inserted the 34x TetO array at three locations in the *Car2* locus. To ascertain which allele the TetO array was inserted on, we used a Cas9-based strategy. For the TetO insertion 4 kb upstream of the *Car2* TSS (EA88), we performed a lipofectamine 2000 transfection of plasmids encoding Cas9 and sgRNAs to cut in the TetO repeats and downstream of the *Car2* open reading frame. For the TetO insertion 107 kb downstream of the *Car2* TSS (EA89), we used sgRNAs targeting TetO and downstream of the Car2 enhancers. To select transfected cells, we added puromycin and hygromycin from 1 day to 3 days after transfection. Five days after transfection, we performed flow cytometry and observed a population of cells with loss of either red or green expression, indicating which allele experienced large deletions and therefore harbored the TetO array. For the TetO array 172 kb downstream of the *Car2* TSS, we used a PCR strategy relying on the existing 10 kb deletion downstream of the *Car2* enhancers on the red allele.

For TACL experiments, cells were transfected with plasmids expressing either TetR-Halo-MAU2 (pAA014) or TetR-Halo (pEA117) using lipofectamine 2000 along with piggyBac transposase plasmid (PB200). Starting two days after transfection, cells were grown in 1 µg/mL puromycin to select for cells with successful integrations. Seven days after transfection, *Car2* reporter expression was measured using flow cytometry. For each location of the TetO integration, two clones were each transfected three separate times. Flow cytometry measurements were normalized by dividing by the average reporter expression in untransfected cells of the corresponding genotype.

### ORCA experiment

#### Cell lines and fixation

ORCA experiments were performed on clonal cell lines. For each TetO insertion position, we subcloned the pool of cells from one TetR-Halo and one TetR-Halo-MAU2 transfection. We measured Halo expression in individual clones by flow cytometry, and then for each condition selected a single clone with a similar expression level. Cells were incubated with 500 nM Halo ligand JFX646 (diluted in media) at 37°C for 30 minutes, excess ligand was removed by incubating the cells in fresh media for 30 additional minutes, and Halo expression was measured by flow cytometry.

For fixation, we dissociated cells using TrypLE, fixed in 4% formaldehyde for 10 minutes at room temperature, washed four times in PBS (by centrifuging for five minutes at 400g), resuspended in 70% ethanol, and stored at -20°C.

#### Multiplexed cell labeling

Cells were labeled by adapting a previously published protocol^50,103^ with minor modifications. Briefly, 3’-amine-modified oligonucleotides (Integrated DNA technologies) were conjugated to Methyltetrazine-NHS ester. The reaction was performed in sodium borate buffer (50 mM, pH 8.5) with DMSO, and the resulting methyltetrazine-oligos were purified by ethanol precipitation. The final products were resuspended in HEPES buffer.

For cell labeling, approximately 2.5-3 million fixed cells were transferred to protein LoBind tubes and 3X washed using 1% BSA in PBS and resuspended in 1X PBS. The cells were first incubated with 1 mM TCO-PEG4-TFP Ester for 5 minutes at room temperature protected from light. Subsequently, the prepared methyltetrazine-oligos were added and incubated for 30 minutes at room temperature. The reaction was quenched using Methyltetrazine-PEG4-Amine and Tris-HCl for 5 minutes. Labeled cells were washed 2X with 1% BSA in PBS and once with 1X PBS only. Cells were resuspended in 70% ethanol in 1X PBS and stored at -20 °C.

#### Cell plating and surface barcode normalization

Barcoded cells were pooled at equal ratios by mixing 2-4 µl of each sample in a single tube. A 10-15 µl aliquot of the pooled cell suspension was then dispensed onto the poly-D-lysine coated central region of a microscope slide and left to air dry for ∼ 5 minutes before proceeding with primary probe hybridization (see below). Multiple slides were prepared in parallel (4-6), and only those with an optimal cell density forming a uniform monolayer were selected for imaging and backup acquisitions. To assess and even out the representation of all conditions prior to the main imaging experiment, barcoded cells were first imaged using readouts targeting the surface barcodes without DNA primary probe hybridization. The relative abundance of each condition was quantified from these barcode signals, and cell numbers were subsequently normalized accordingly.

#### Probe design

The *Car2* locus (chr3:14,753,359-15,369,435) was tiled with 72 barcodes spanning 8 kb each, with every barcode targeted by ∼120 individual probes. Each probe was 100 bp in length and compromised of 4 segments: 20 bp of a reverse and forward primer sequences used during probe synthesis, 20 bp complementary to the fiducial probes, 20 bp of barcode sequence, and 40 bp of unique genomic targeting sequence. Probes were designed using previously described software^104^, which is accessible at https://github.com/BoettigerLab/ORCA-public. The probe library was synthesized as an oligonucleotide pool by GenScript.

#### DNA hybridization

Hybridization with DNA primary probes was carried out following^105^. Briefly, following pooling and plating, cells were fixed in 4% paraformaldehyde (PFA) for 10 minutes and washed 3X with 1X PBS, which was gently added to the slide by pouring along the side of the 6 cm Petri dish to avoid direct flow over the cells. Cells were then premetallized in 20-40 µl of 0.5% Triton X-100 in 1X PBS for 10 minutes and rinsed 3X with 1X PBS. Permeabilized cells were then incubated in 20-40 µl 0.1 M HCl for 5 minutes and washed 3X with 1X PBS. To remove RNA, cells were treated with 20-40 µl RNase A (10 µg ml ^-1^) for 30 minutes at 37 °C and then washed 3X with 2X SSC. Next, cells were equilibrated for 35 minutes in hybridization buffer (2X SSC, 50% formamide, 0.1% Tween-20). The volume of solution applied varied depending on the area occupied by cells on each slide; solutions were added in sufficient volume to ensure complete coverage of all cells.

Hybridization mixture was prepared by mixing 2 µl of 840 ng µl^-1^ of DNA primary probes and 30 µl hybridization solution (2X SSC, 50% formamide, 10% dextran sulfate, and 0.1% Tween-20). The mixture was then applied to the cells, covered with a coverslip treated with Sigmacote, denatured for 3 minutes at 90 °C and then incubated overnight in a humidified chamber at 42 °C. The following day, samples were 3X washed with 2X SSC and post-fix in a mixture of 2% glutaraldehyde (GA) and 8% PFA in 1X PBS for 30 minutes. After post-fixation, slides were 3X washed with 2X SSC and mounted on the microscope for sequential labeling and imaging, starting with surface barcode imaging.

#### ORCA Imaging setup

Imaging was performed using a custom-built ORCA microscope setup with the complete configuration parameters available via MicroMeta App^106^. Briefly, the system was constructed around a Nikon Ti2 body with a large 32 mm aperture optics enabling the acquisition of wide-field images (2,048 x 2,048-pixel area with a final pixel size of 108 nm). Illumination was provided by solid-state laser arrays by Lumencor (CELESTA light engine) paired with a custom polychroic mirror (Chroma, ZET408/473/545/635/750rpc-uf2) and emission filter (Chroma, ZET408/473/545/635/750m). The system was equipped with a Marzhauser Wetzlar xy-stage (00-24-625-000) and a Mad City Labs Inc. Z objective positioner (Nano-F200S) for z-axis control. The setup was completed with a Hamamatsu ORCA-Flash4.0 V3 sCMOS camera, an IR-laser-based autofocus system, and custom fluidics set up^105^.

#### ORCA image acquisition

Primary probe hybridized samples were imaged by detecting readout sequences with dye-labeled secondary oligonucleotides: AlexaFluor 750 for DNA readouts, Cy5 for surface barcodes and Cy3 for fiducial markers, with the Cy3 channel used for accurate spatial registration and sample drift correction. Imaging was started with surface barcodes and followed by DNA tracing. Fiducial signals were detected during the first DNA imaging round together with the first DNA readout and were subsequently imaged in every following cycle.

For each readout barcode, ∼ 400 µl of probe solution (25% ethylene carbonate in 2× SSC containing the readout oligonucleotide and displacement oligos complementary to the previous readout) was flowed into the chamber and incubated for 30 minutes to allow hybridization and strand displacement. All remaining wash, buffer exchange, and acquisition steps were performed as in the published protocol^105^.

#### ORCA analysis

Raw imaging processing was performed following the image processing pipeline detailed in ref^105^ with published scripts available at: https://github.com/BoettigerLab/ORCA-public. This pipeline yields 3D coordinates for every barcode within a given trace, which were subsequently transformed into pairwise distance matrices for structural analysis.

#### Computational correction for underperforming readouts

To ensure the integrity of population-level data, we computationally corrected all median distance and contact frequency maps for artifacts introduced by specific readout probes. For most of these readouts, the poor performance stems from flawed intrinsic properties, a recurring issue documented in prior ORCA experiments. In this study, a specific list of seven readouts (5, 10, 16, 21, 23, 48, and 65) were identified as unreliable based on visual inspection of the raw imaging data, which revealed issues such as weak signal or spurious fluorescence. The data corresponding to these specific rounds was masked and the resulting gaps in the data matrices were filled by interpolating values from neighboring, high-quality hybridization points.

Readout 38 was masked to account for the 10 kb deletion in the TetO cell lines. While the readout itself performed well, this step prevents the deletion from creating a strong predicted artifact in the differential contact and distance maps for TetO vs. non-TetO conditions.

#### Demultiplexing cell surface barcodes

Demultiplexing was performed using a custom MATLAB script that integrates Cellpose for nuclei segmentation^107^. For each field of view (FOV), a maximum intensity projection image was generated from the drift-corrected barcode imaging channels to facilitate robust segmentation. Cellpose was then run with its pre-trained ‘nuclei’ model to generate segmentation masks. For each segmented nucleus, a barcode intensity vector was created by calculating the median pixel intensity within the mask for each barcode channel. These vectors were normalized to account for variations in probe brightness. Finally, each nucleus was assigned an identity by matching its normalized barcode vector to a predefined codebook using a k-nearest neighbor approach, with an associated confidence score calculated to assess assignment ambiguity. The resulting table linked each identified spot from the primary experiment to its parent nucleus and decoded identity.

#### Allele disambiguation

To distinguish between control allele (non-TetO allele) and allele engineered for forced cohesin loading (TetO allele) via a 10 kb deletion, a custom Python script was used to classify chromosomal traces in each demultiplexed cell based on the presence or absence of imaging readout 38, which specifically hybridizes to the deleted region in the *Car2* control allele. To ensure high confidence in our classification, this comparison was performed exclusively between the two alleles within the same nucleus, which serves as an internal control to mitigate potential misalignment errors or technical false negatives arising from variable hybridization efficiency.

#### Quantification of enhancer-promoter interactions

Enhancer-promoter interactions were assessed between windows of loci centered around the *Car2* enhancers and promoter. Specifically, we used a +/- 1 bin window for each (P-window, E-window) to account for the physical span of the regulatory elements and to mitigate potential signal dropout from suboptimal probe detection efficiency during imaging.

To avoid using an arbitrary distance cutoff, a data-driven contact threshold was determined as the median of all pairwise distances between adjacent probes measured across the entire traced *Car2* domain in the control cell population, approximately 200 nm across conditions. An E-P interaction was then classified “in contact” for a given cell if the minimum distance between the E and P windows was less than or equal to this threshold.

To capture the full spectrum of interaction distances without binarization, we analyzed the distribution of minimum E-P distances. For each individual cell, we identified the single minimum distance between any locus in the E-window and any locus in the P-window. This metric represents the closest physical approach between the two regulatory regions within that cell.

### Simulations

### 1D lattice model

We performed one-dimensional lattice simulations to model loop extrusion dynamics. Loop extrusion parameters were chosen to reflect mESC dynamics, and the various loading and barrier configurations below were used to probe different determinants of cohesin accumulation. We modeled 1D dynamics with fixed-timestep simulations on a lattice where each lattice site represented 250 base pairs of genomic sequence. All lattice sites shared a uniform unloading (or dissociation) rate for extruders; this is specified via the processivity (330 kb, equivalent to 1320 lattice sites or 660 lattice sites for each arm), the average distance extruded before dissociation. Each site could either represent a generic genomic location with baseline extruder loading probability or be designated as a boosted loading site with increased extruder loading frequency. For standard sites, cohesin loading occurred with uniform probability across the lattice. Average spacing between extruders was determined by a separation parameter, set to 185 kb (740 lattice sites), following experimental estimates^108^ and previous simulations^23^ in mESC.

Any extruder that dissociated from the lattice was immediately reloaded at a random unoccupied lattice site, where the likelihood of reloading at a given position was proportional to its local loading rate. Each extruder occupied two adjacent lattice sites, corresponding to its left and right legs. At each discrete time step, extruder legs not stalled by a barrier could move outward, (i.e. the left leg moves left and the right leg moves right). For example, an extruder can update from position (i, j) to (i–1, j+1). Extruder legs could not bypass one another upon collision and could not exit the first or last lattice sites, effectively confining all movement within the lattice boundaries. Positions of extruders were recorded after discarding the initial 20% of simulation steps to compute observables.

Dynamic CTCF barriers were implemented as unidirectional obstacles that selectively stalled extruder legs based on their direction of motion (as previously^45^). Dynamic barrier occupancy was modeled as a two-state process, governed by defined binding and unbinding times. Extruders encountering an unbound barrier site continued extrusion uninterrupted. An extruder leg moving in the direction opposite to the orientation of an occupied barrier became stalled upon contact. While one leg was stalled at a bound barrier, the other leg was allowed to continue moving independently.

#### Boosted loading site layouts

Boosted loading was reported as a fold-increase per kilobase. E.g., a loading boost of 100 on a 250 bp site is equivalent to a fold-increase of 25 per 1 kb. In simulations involving a single focal loading site, boosted loading was applied at a single 250 bp lattice site. For simulations representing larger loading regions or “active TADs” of uniformly boosted loading, multiple adjacent boosted loading sites were used. In all scenarios, boosted loading added to the total number of extruders in the simulated region. One boosted loading region or site was considered per 1,000 lattice sites. We averaged over 30,000,000 simulation snapshots for these layouts.

#### Quantifying cohesin at the loading site

To quantify the relative enrichment of extruders at boosted target loading sites (Fig. 3B and Extended Data Fig. 5C), we counted extruder reads within a four-site window (corresponding to 1 kb) centered on the center of each boosted target loading region. This local read count was divided by the average number of extruder reads measured in two distal control regions located 20 kb to the left and 20 kb to the right of the target region, using windows of the same size. This value was then normalized to the corresponding value obtained for the uniform loading scenario. The resulting ratio quantifies local extruder enrichment at the boosted loading site while controlling for local background read density.

#### Boosted loading between inward-oriented CTCF layout

To clarify the impact of targeted loading on accumulation at nearby CTCFs, we considered simplified scenarios with isolated CTCF barrier sites oriented inwards symmetrically at a defined distance from a loading site (Fig. 3D, Extended Data Fig. 5). Such arrangements were placed at the center of a 1,000 site region and tiled ten times to form a 10,000-site lattice. For these simplified scenarios, we set the CTCF bound time to be approximately one quarter of the cohesin residence time, reflecting previous estimates^45^. Unless noted, isolated barriers had an occupancy of 0.5, meaning they were CTCF-bound 50% of the time and unbound the other 50%. We calculated an enrichment ratio at CTCF sites as the accumulation of extruder legs at barrier sites in a targeted loading scenario normalized by the baseline from uniform loading. We computed in silico ChIP-seq profiles by recording the positions of extruder legs from 30,000,000 instances of 1D lattice simulations for this layout.

#### Tandem CTCF layout

To precisely quantify the impact of nearby CTCF sites arranged in tandem (Fig. 6D), we performed a set of simulations with otherwise isolated tandemly-oriented pairs of CTCF sites. A set of tandem sites were placed at the center of a 1,000 site region and tiled ten times to form a 10,000 site lattice. We quantified how the front CTCF can block accumulation at the back CTCF by varying the occupancy of the front CTCF while keeping the back CTCF at 0.5 occupancy. We then calculated RAD21 reads normalized by the number of times each barrier appeared at its genomic position (RAD21/CTCF). We computed in silico ChIP-seq profiles by sampling extruder leg positions from 100,000,000 1D lattice simulation instances.

#### Variable spacing and occupancy of CTCF barriers layouts

To develop simulations with more complex patterns of cohesin accumulation closer to those in cells, we designed simulations with randomly distributed barriers with variable bound times and occupancies as follows (Fig. 6E, Extended Data Fig. 10) for a lattice with 300,000 sites. Barrier occupancies were independently sampled with replacement from empirical measurements of CTCF binding via single-molecule footprinting^109^, and the total number of barriers (multiplied by their occupancy) was chosen to match experimental estimates. Specifically, we sampled barriers until their combined occupancy reached 133 in 10 Mb, based on estimates for the number of bound CTCF molecules in a region of this size (∼217,000 total CTCF molecules in a mESC nucleus, with 50% bound (based on FRAP measurements^110^), and an average estimated genomic content per cell ∼3 × 2.7 Gb^108^). Lattice positions and orientations were randomly assigned for each sampled barrier. While per site occupancies are available experimentally, per site bound and unbound times are not. Based on previous estimates^45^ we thus set these dynamic parameters by scaling sampled bound times such that strong sampled barriers (0.65 occupancy) had a residence time (656 sec) of roughly one third the extruder lifetime, thus fixing their unbound time (353 sec). Average sampled barriers (∼.35 occupancy) had a correspondingly lower residence time (233 sec). For varying bound time simulations, we sampled site occupancies, and determined corresponding site bound times using the fixed unbound time (353 sec). We then introduced variability in bound times at fixed sampled occupancies by: drawing an updated bound time from a Gaussian distribution with mean equal to the sampled bound time and standard deviation of ⅓ of the sampled bound time and rescaling the unbound time to maintain the sampled occupancy. For the varying unbound time simulations, a similar rescaling procedure was used, albeit determining site unbound times from the sampled occupancy and a fixed bound time (233 sec). For constant occupancy simulations, all barriers had the same occupancy, and thus the same bound (233 sec) and unbound times (353 sec). For each scenario, we ran 100 independent simulations using multiprocessing, with approximately 153,000 recorded instances per simulation. Statistics were computed by aggregating data across all runs, yielding a total of approximately 15,300,000 simulated instances.

To evaluate the effect of variable separation between barriers on extruder accumulation, we conducted simulations using regular barrier arrangements. First, we maintained the same barrier density (per 10 Mb) and positioned barriers at constant intervals of ∼ 35 kb with alternating orientations (right, left, right, etc.). To assess the impact of barrier spacing on the dispersion of cohesin accumulation, we also increased the distance between barriers six-fold.

To quantify cohesin enrichment, we calculated the accumulation of extruders at barrier sites. To compare simulations with experimental data, we scaled both RAD21 and CTCF read counts to match the read densities measured by ChIP-seq. Specifically, we calculated the total number of simulated extruder and CTCF reads per base pair within the 75 Mb simulated region, and applied a normalization factor such that the simulated read density equaled the experimental read density (0.016764 reads per basepair, per haploid genome). This normalization ensured that simulated and experimental datasets had comparable read coverage per genomic length.

To further quantify the role of barrier spacing, we analyzed simulated cohesin positioning at barriers similarly to experimental data. To investigate the role of nearby CTCF sites in pairs oriented tandemly), we further calculated RAD21 reads normalized by how many times each barrier appeared at its genomic position (RAD21/CTCF). This normalization allowed us to measure RAD21 accumulation specific to each individual barrier. Specifically, we identified pairs of tandemly oriented barriers, and compared the relative enrichment of extruders between them by computing the ratio of extruder accumulation in the first versus the second barrier, stratified by their spacing. Then we computed the cumulative distribution functions over their spacing separately for right-oriented and left-oriented barriers (Extended Data Fig. 11B). Finally, we assessed how extruder accumulation (for all orientations) scales with barrier proximity by comparing extruders versus barriers read counts across random configurations. We binned barriers by their nearest-neighbor distance and calculated the average extruder/barrier ratio in each bin (Extended Data Fig. 11A).

#### 3D polymer simulations

We performed 3D polymer simulations for the targeted loading scenarios by modeling a 2.5 Mb segment of chromatin as a 50 nm polymer fiber, with each monomer corresponding to 2.5 kb of genomic DNA or 10 sites in the corresponding 1D lattice model. To implement the 3D polymer simulations, we used the Polychrom framework (https://github.com/open2c/polychrom), which leverages OpenMM^111^ to simulate large ensembles of chromatin conformations. To simulate loop extrusion, we performed 200 3D steps between each 1D lattice update. Based on previous timescale matching^45^, each simulated time step in 3D represents δt_3d_ = 0.015 sec to give a mean extruder residence time on chromatin of approximately 33 minutes. Loop extrusion was modeled by introducing harmonic bonds between the two monomers representing the arms of each extruder (positions *I* and *j*). As the loop extrusion factor progressed, these bonds were dynamically updated; removing the bond between *(I, j)* and adding a new bond between *(i–1, j+1)* to reflect loop enlargement. Under wild-type conditions, the polymer exhibited Rouse dynamics.

#### In silico Hi-C analysis

For in silico Hi-C analysis, we generated 3,000,000 polymer conformations from our 3D simulations for every parameter set we considered. Using these conformations, we constructed Hi-C contact maps at a resolution corresponding to the size of one monomer (2.5 kb), applying the default *polychrom* (https://github.com/open2c/polychrom) capture radius of 2.3 monomers to define contact events. This threshold closely matches empirical contact distance estimates^112^. Simulated contact matrices were converted into .cool format using *polykit* (https://github.com/open2c/polykit), facilitating analysis within the standard Hi-C data processing framework^113,114^.

#### Code

Code specifying simulations and for performing analysis are available at: https://github.com/Fudenberg-Research-Group/Targeted_Cohesin_Loading

### Determining the fraction of cohesin reads at CTCF sites

To measure the fraction of cohesin reads at CTCF sites, we first performed calibrated ChIP-seq with RAD21 (abcam ab154769) and SMC1 (Bethyl A300-055A) antibodies in untagged (E14) cells, untreated RAD21-eGFP-AID (EN272.2) cells, and EN272.2 cells in which RAD21 was degraded by adding 500 mM of Indole-3-Acetic Acid sodium salt (IAA, auxin analog, Tocris Bioscience 7932) to the culture media for 3.5 hours. The position of the degron tag precluded use of the Abcam ab992 RAD21 antibody used in other experiments. For each of two replicates, the number of cohesin reads in CTCF peaks was computed as in https://github.com/Fudenberg-Research-Group/fastaFRiP using deeptools.countReadsPerBin and then multiplied by the scaling factor. The background-corrected total number of reads was calculated by (total mouse reads * scaling factor) – (total reads after RAD21 depletion * scaling factor). Finally, the corrected fraction of reads in CTCF peaks was calculated by (cohesin reads in CTCF peaks) / (background-corrected total reads).

### Analysis of cohesin/CTCF ratios

We performed four replicates of calibrated ChIP-seq with CTCF (Active motif catalog number 61311) and RAD21 (Abcam ab992) antibodies in E14 ESCs. In NPCs, we performed two replicates each of CTCF and RAD21 ChIP-seq. For each protein, we combined equal numbers of reads from each of the ChIP-seq replicates. Approximately 46 million total reads were included for CTCF and RAD21 in ESC. For NPC analysis, the number of reads was normalized to match the number in ESCs. We used bedtools window to calculate the number of reads in each CTCF peak.

To calculate correlations in the number of reads in CTCF sites between ChIP-seq datasets, we used multiBigwigSummary and plotCorrelation from deeptools with the Pearson correlation method.

For comparing cohesin/CTCF ratios to the distance to the nearest enhancer, we used the enhancer list from^39^. For transcription start sites, we used transcript coordinates from GENCODE VM23 and included protein-coding genes with at least 1 transcript per million. Gene expression levels are from ENCODE RNA-seq in E14 mouse ESCs (identifier ENCSR000CWC)^115^. A and B compartments were called using the compartment_score function in GENOVA^116^ with Hi-C data from two combined replicates of wild-type mESCs^23^.

For the analysis of CTCF motif variants (Extended Data Fig. 6F), we used core, forward, and reverse motifs from^53^. We used FIMO from the MEME suite^117^ to identify instances of the motifs (p < 0.0001) in the CTCF peaks. We plotted the RAD21/CTCF ratios for peaks in the following categories: only core motifs; at least one core and at least one upstream motif but no downstream motif; at least one core and at least one downstream motif but no upstream motif; no core motifs.

For analysis the of the distance over which CTCF sites can block cohesin traffic, we counted the number of forward and reverse core CTCF motifs^53^ in each CTCF peak. We selected pairs of peaks that each contained only forward motifs and for which the next downstream peak was over 100 kb away, and similarly selected pairs of peaks with reverse motifs. For each pair, we calculated log_2_[(first peak RAD21/CTCF)/(second peak RAD21/CTCF)] and plotted the cumulative distribution of this metric based on the distance between the peaks.

### Micro-C analysis

We analyzed micro-C from^118^, using data with two pooled replicates from untreated mESCs (GSE178982_CTCF-UT_pool.mcool) or cells with 3 hours of CTCF depletion (GSE178982_CTCF-AID_pool.mcool). Plots show data normalized by the expected interaction frequencies. We ranked MAU2 peaks based on the number of ChIP-seq reads, and generated a pileup at the top one third of sites using cooltools.pileup^114^. Control sites were selected using bedtools shuffle^119^ to choose random sites within TADs that contain a top MAU2 site, with TAD definitions from^118^. Repeated permutations of random sites yielded consistent pile-up plots.

### Enformer

As Enformer is not trained on relevant mouse ESC datasets, we generated predictions for cohesin and CTCF binding in mouse CH12 cells. We downloaded CH12 CTCF and RAD21 ChIP-seq data from the ENCODE portal (identifiers ENCSR000ERM and ENCSR000ERK), mapped the reads as above, and called peaks using macs2. We selected a CTCF peak with a low RAD21/CTCF ratio (chr15:97665011-97665743) to use as a recipient locus, and selected ten CTCF peaks with higher CTCF/cohesin ratios to use as donor sequences. We then generated test sequences by swapping 512 bp (centered at the center of the CTCF peak) from the donor sequence into the recipient sequence. We used Enformer^120^ to predict CH12 RAD21 and CTCF binding at the test sequences. We plotted the sum of the cohesin signal across the 512 bp divided by the sum of the CTCF signal across the 512 bp. We then modified the recipient sequence by replacing the CTCF motifs in the CTCF peaks flanking the central peak with neutral sequence (replacing 17 bp at each of four motifs), and repeated the swapping of the donor sequences into the modified recipient sequence.

## Supporting information

targeted loading extrusion movie

Plasmids_cell-lines_sgRNAs

## Acknowledgements

We are grateful to members of Daniele Canzio’s group for discussion and access to equipment, and to Alex Buckley, Simon Gaudin, Fernanda Vargas-Romero, Elzo de Wit, and Luca Giorgetti for critical feedback on the manuscript. We thank David Spector and Stanley Qi for the U2OS 2-6-3 cell line. Funding sources: Grant NIH R35GM142792 to EN, Grant NIH R35GM143116 to GF, the Chan-Zuckerberg Biohub San Francisco Investigator program to EN, the Sandler Program for Breakthrough Biomedical Research at UCSF, NSF EAGER grant 2313792 to EN, NIH training grant T32HL007731 (ECA), fellowships from the Helen Hay Whitney foundation (ECA), the UCSF-California Institute for Regenerative Medicine Scholars Training Program education grant EDUC4-12812 (ECA and KH), and KAUST Ibn Rushd Fellowship (AA). Part of the sequencing was performed at the UCSF CAT, supported by UCSF PBBR, RRP IMIA, and NIH 1S10OD028511-01 grants. Flow cytometry was performed in the Laboratory for Cell Analysis in the UCSF Helen Diller Family Comprehensive Cancer Center, supported by grant P30CA082103.

## Author contributions

ECA performed, analyzed, or supervised all experiments except ORCA, with technical support from EMA, ASA, RS, KH, and EPN. HR performed simulations under the supervision of GF. AA designed, performed and analyzed ORCA with support from IC and under the supervision of ANB. EP designed the study with ECA and input from GF. ECA assembled figures and wrote the manuscript with assistance from EPN, GF, and input from all co-authors.

## Ethics declaration

No competing interests to declare.

## Data availability

Sequencing data was deposited to GEO GSE278338.

## Code availability

https://github.com/hrahmanin/Target_cohesin_loading

## Extended Data Figures

**Extended Data Figure 1:**
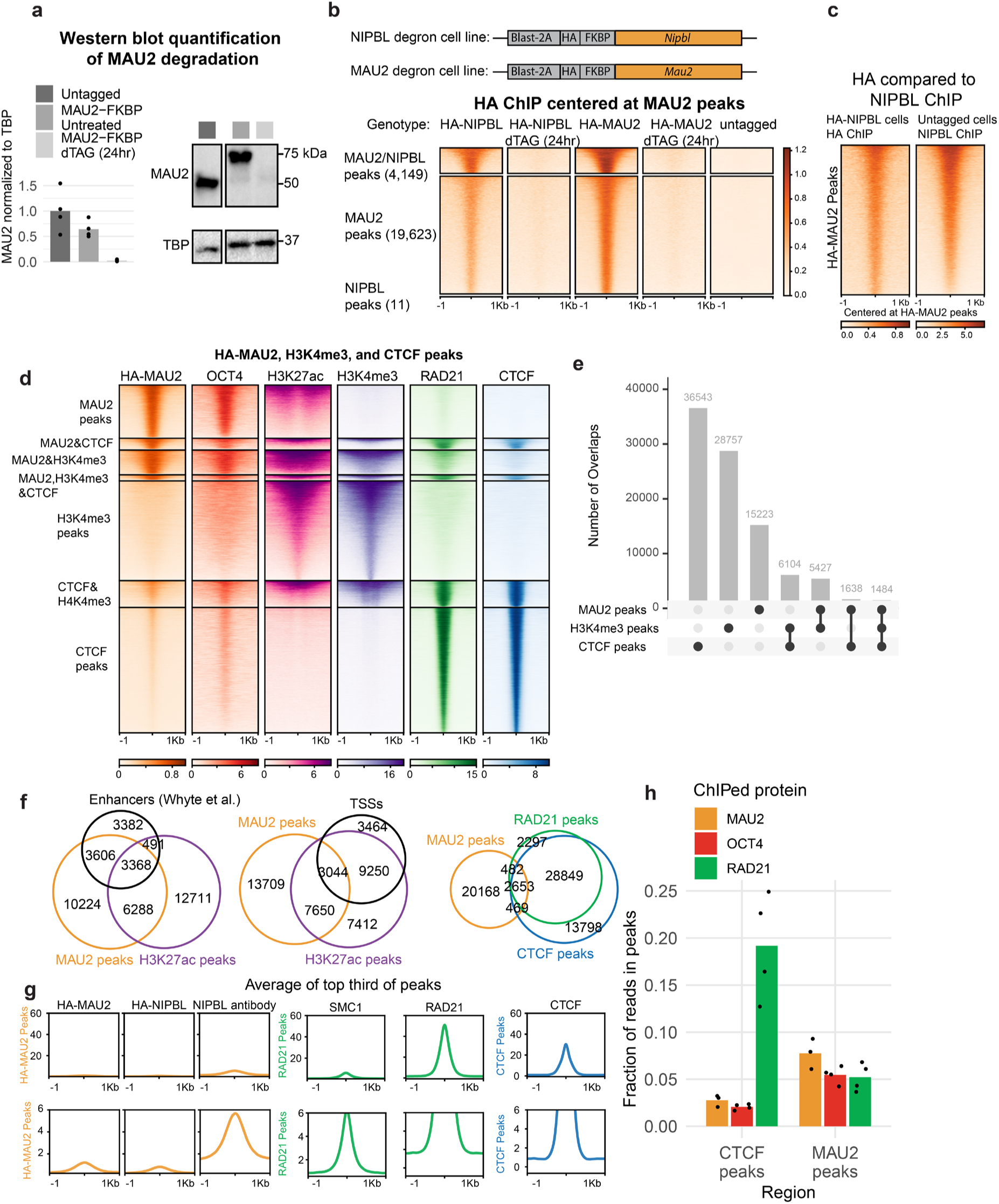
Reproducible NIPBL/MAU2 binding sites overlap with enhancers in mouse ESCs. a, Western blots show depletion of MAU2-FKBP after 24 hr dTAG treatment. **b**, HA ChIP-seq signal at MAU2 and NIPBL peaks is lost after NIPBL or MAU2 degradation (24 hr dTAG treatment). NIPBL and MAU2 co-bind the same sites, with the more sensitive HA-MAU2 ChIP-seq detecting more peaks. **c**, HA-NIPBL ChIP-seq signal closely matches ChIP-seq with an endogenous NIPBL antibody. **d**, Integration of ChIP-seq datasets highlighting that MAU2 binds the subset of active chromatin sites (H3K27ac) that bind OCT4 (enhancers). RAD21 (cohesin) is enriched at CTCF sites, with minimal signal at enhancers/MAU2 sites. OCT4 data is from ref^51^ and H3K4me3 from ref^115^. **e**, Upset plot showing that MAU2, H3K4me3 (marking promoters), and CTCF largely bind distinct loci. **f**, Overlap of MAU2 peaks with active chromatin (H3K27ac peaks), enhancers (defined as sites bound by pluripotency transcription factors OCT4, SOX2, and NANOG from ref^39^), TSS regions (defined as 1 kb upstream of TSSs of GENCODE protein-coding genes expressed at >1 transcript per million), RAD21 peaks, and CTCF peaks. **g**, Average binding profiles of NIPBL/MAU2, cohesin subunits, and CTCF at their own peaks, showing that NIPBL/MAU2 binding sites have weak enrichment over background. Plots are shown on two different scales for comparison. **h**, Fraction of MAU2, OCT4, and CTCF ChIP-seq reads found in either CTCF or MAU2 peaks. OCT4 ChIP-seq is from ref^51,121^.

**Extended Data Figure 2:**
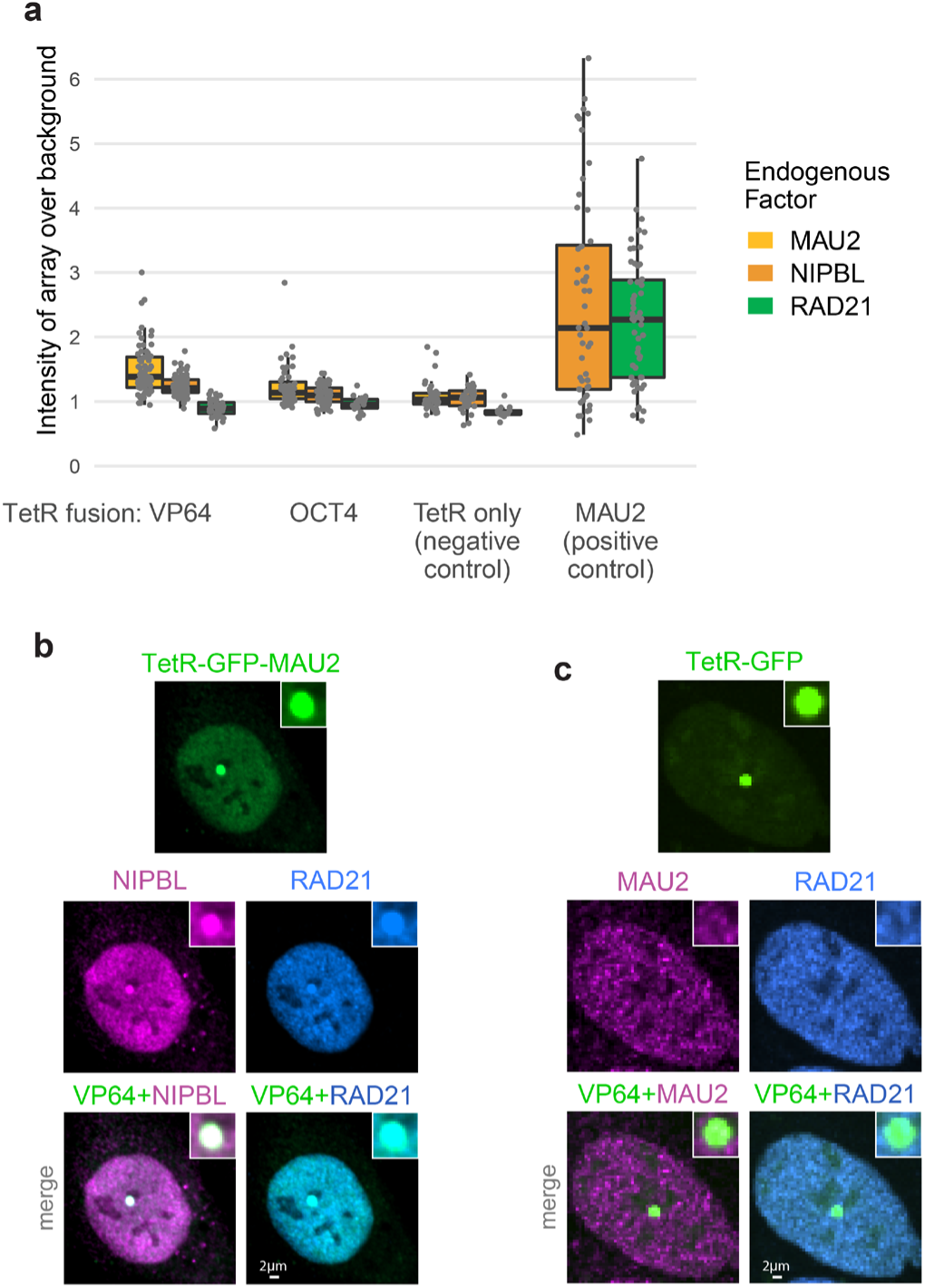
Recruitment of cohesin by TetR-MAU2. **a**, Intensity of MAU2, NIPBL, or RAD21 immunofluorescence signal at the TetO array divided by intensity of the background nuclear signal. **b**, Example cell showing that TetR-MAU2 recruits NIPBL and RAD21 to the TetO array. **c**, Example cell showing that TetR alone does not recruit MAU2 or RAD21 to the TetO array.

**Extended Data Figure 3:**
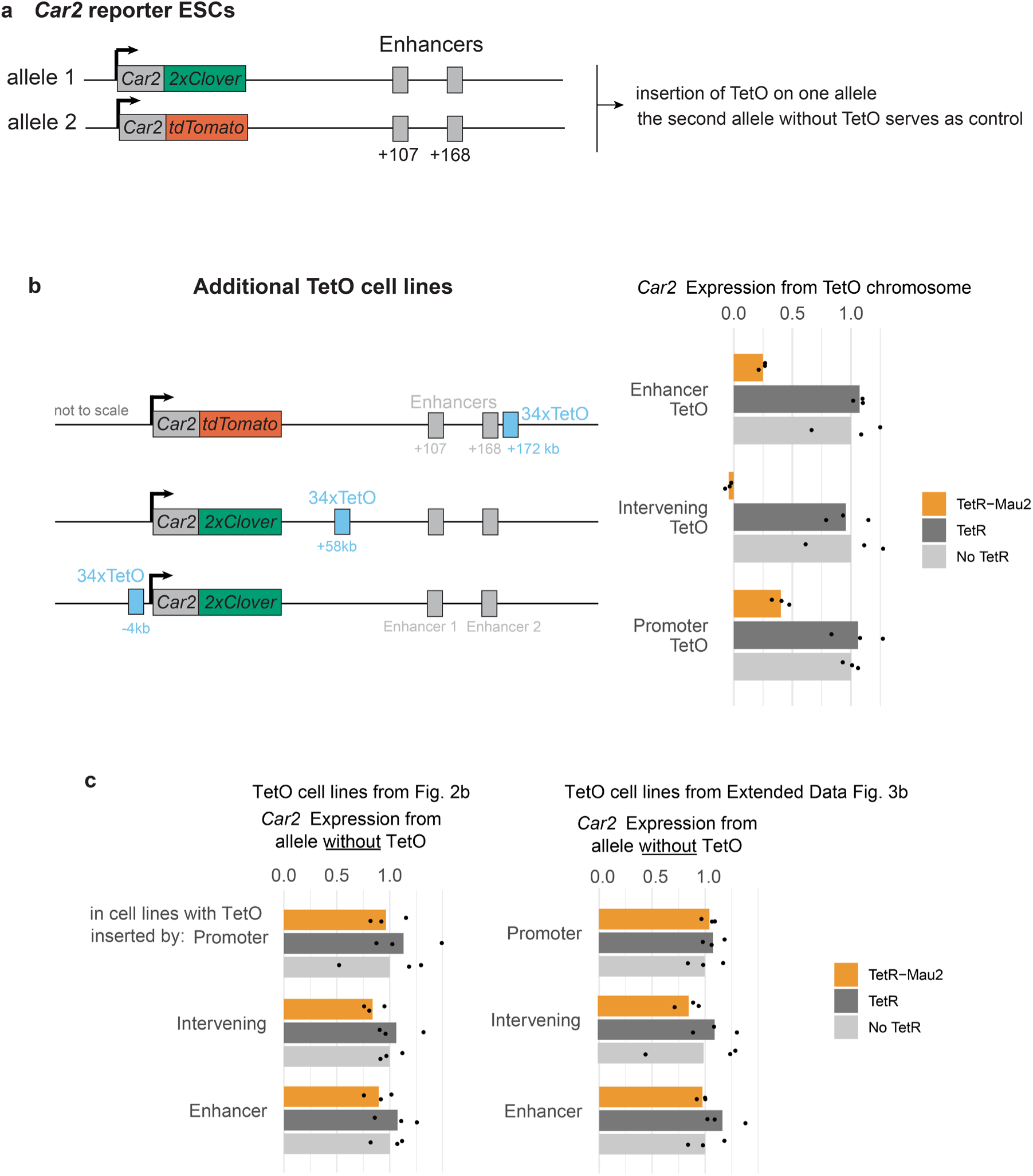
Additional clones for targeted cohesin recruitment. **a**, *Car2* reporter ESCs (from ref^42^): in each cell, one *Car2* allele harbors a tdClover reporter and the second allele harbors a tdTomato reporter. *Car2* expression in cells with heterozygous TetO insertions can therefore be directly compared between the TetO allele and control allele without the TetO. **b,** Additional cell lines with TetO insertions either on the tdClover or the tdTomato allele, replicating observations reported in Fig. 2. **c**, In the same cells that displayed reduced *Car2* expression on the TetO allele upon TetR-Mau2 expression, *Car2* expression from the control allele without the TetO insertion remains unchanged across three biological replicates from separate transfections.

**Extended Data Figure 4:**
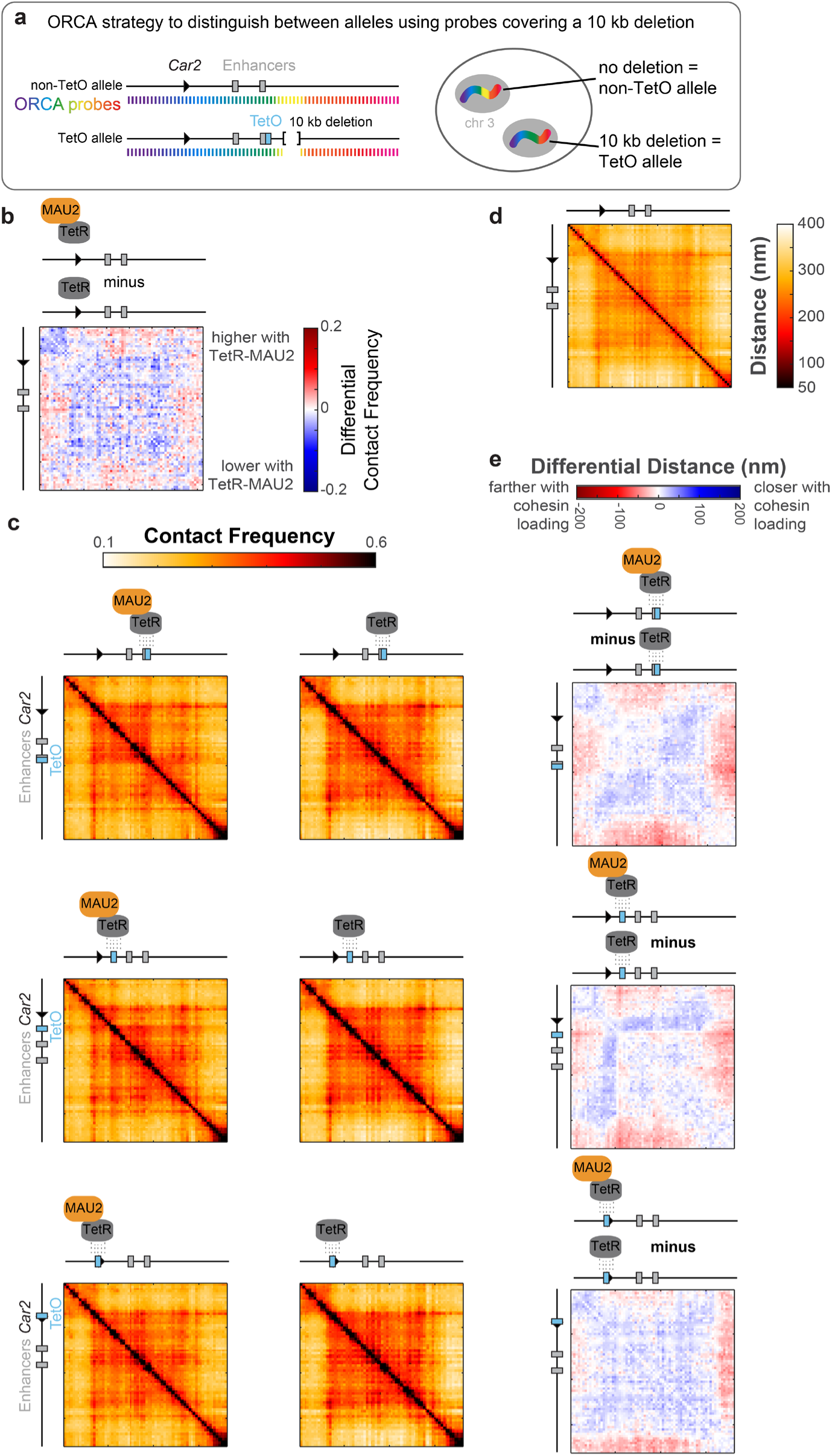
ORCA measurements comparing alleles with and without high cohesin loading. **a**, The TetO array used for cohesin recruitment is inserted on one allele, where we also engineered a neutral 10 kb deletion to distinguish the two alleles. This deletion enables distinguishing the two alleles, as the TetO allele will not display ORCA signal from probes covering the 10 kb deletion. **b**, ORCA interaction frequencies from the non-TetO allele are reduced upon TetR-MAU2 overexpression compared to cells expressing TetR alone. This comparison indicates that MAU2 expression slightly diminishes ORCA interaction frequencies. **c**, ORCA contact frequencies (∼200 nm, see Methods) for the TetO allele in cells expressing TetR-MAU2 or TetR only (8 kb resolution). **d**, Heatmap showing the average distances at the *Car2* locus. **e**, Subtraction heatmaps comparing average distances in cells expressing TetR-MAU2 or TetR alone.

**Extended Data Figure 5:**
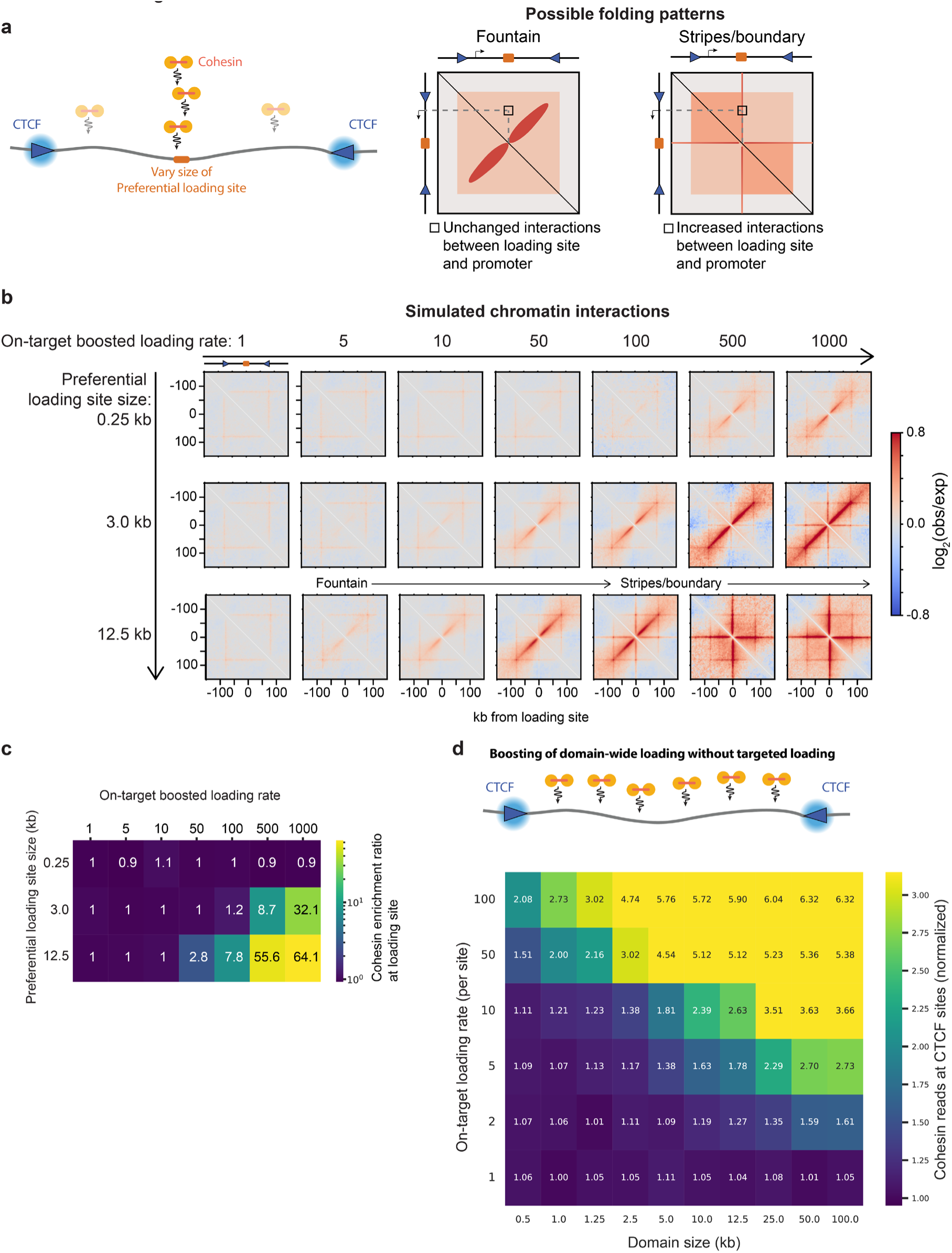
Width of cohesin loading site affects cohesin blocking and traffic. **a**, 3D polymer simulations in which the rate of cohesin loading is boosted at a loading site of varying size. **b**, Simulated Hi-C-like contact maps (2.5 kb resolution). Increasing cohesin loading creates fountain structures. For large loading sites with high loading, stripe patterns also emerge from the loading site, as colliding cohesin molecules block each other. **c**, 1D lattice simulations as in Figure 3A and 3B show that cohesin accumulates at loading sites only if loading is high across multiple adjacent monomers. The enrichment ratio at the loading site is calculated as the amount of cohesin in 1 kb at the center of the loading site divided by the amount of cohesin in 1 kb at a background site. **d**, Simulation exploring how boosting cohesin loading across domains of increasing size affects cohesin abundance at flanking CTCF sites.

**Extended Data Figure 6:**
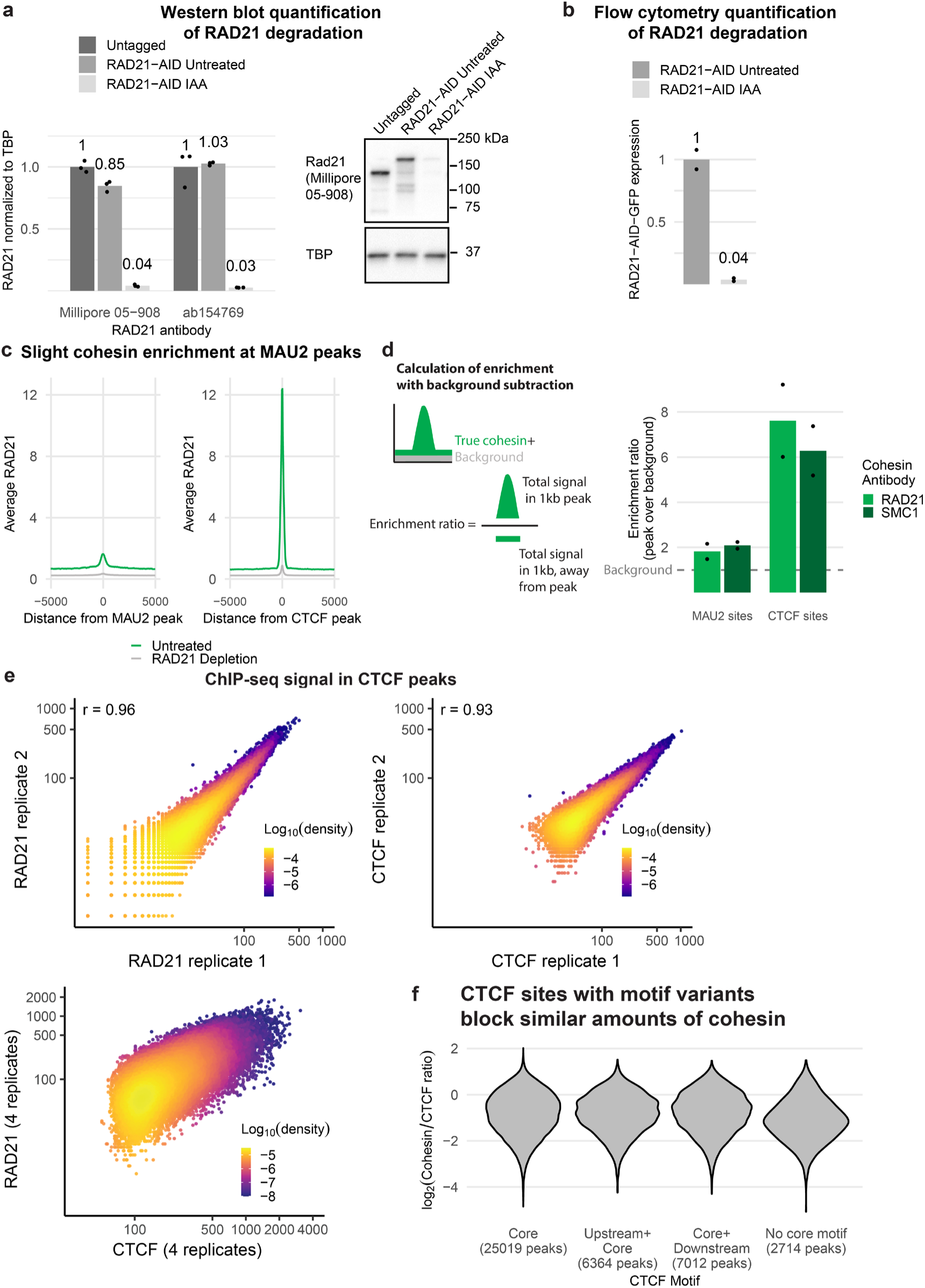
Cohesin is slightly enriched at MAU2 peaks, and cohesin at CTCF peaks is weakly correlated with CTCF occupancy. **a**, Western blots and **b**, flow cytometry show that in RAD21-AID-GFP cells treated with IAA for 3.5 hours, >95% of RAD21 was degraded. **c**, Average RAD21 signal at MAU2 peaks (that do not overlap CTCF peaks) or at CTCF peaks before or after 3.5 hr RAD21 depletion. **d**, Average enrichment of cohesin binding at MAU2 or CTCF peaks after background subtraction. **e**, At CTCF peaks, the number of reads is highly correlated between replicates of CTCF or RAD21 ChIP-seq but shows wider dispersion between RAD21 and CTCF (four combined replicates reproduced from Fig. 5B for comparison). **f**, RAD21/CTCF ratios for CTCF sites based on the presence of upstream or downstream CTCF motifs^53^. Sites with upstream, downstream, or only core motifs show similar distributions, while peaks with no identifiable core CTCF motif block less cohesin.

**Extended Data Figure 7:**
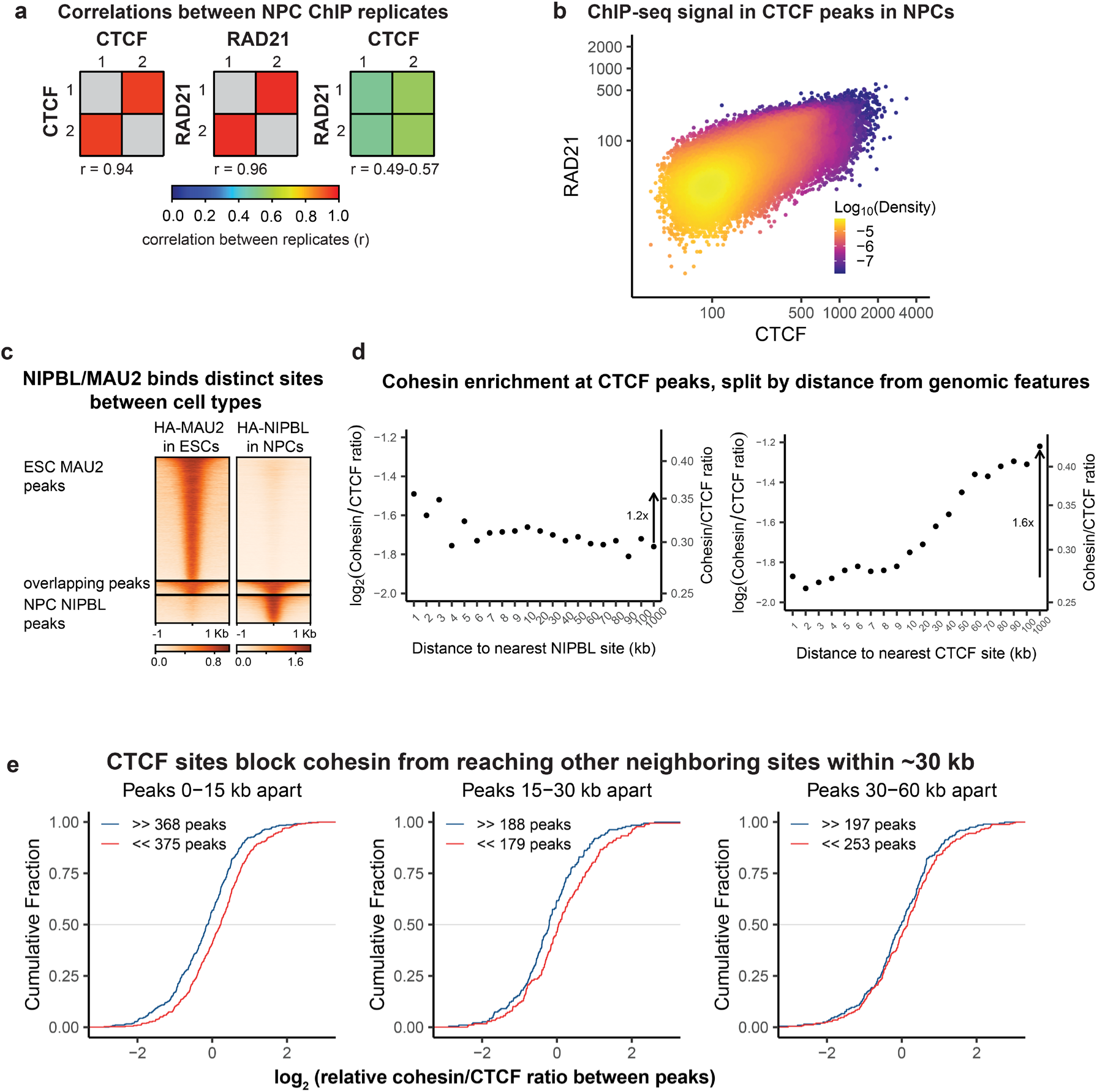
CTCF sites, more than NIPBL/MAU2 sites, control cohesin traffic in Neural Progenitor Cells (NPCs) differentiated from ESCs. **a**, At CTCF peaks, the number of ChIP-seq reads is more highly correlated between two CTCF replicates or two RAD21 replicates, compared to the correlation between CTCF and RAD21 (Pearson correlation coefficient). **b**, RAD21 vs. CTCF reads in each CTCF peak. Read numbers were normalized to match the number in ESCs in Fig. 5B. **c**, NIPBL/MAU2 binding sites are largely not overlapping between ESCs and NPCs. **d**, Average ratio of cohesin/CTCF reads in CTCF peaks based on the distance from the peak to the nearest NIPBL or CTCF site. **e**, The distance over which CTCF sites block cohesin in NPCs was analyzed as in Fig. 6C. Similarly to ESCs, pairs of CTCF sites with forward motifs have lower relative cohesin/CTCF ratios than pairs with reverse motifs only when the peaks are within 30 kb of each other.

**Extended Data Figure 8:**
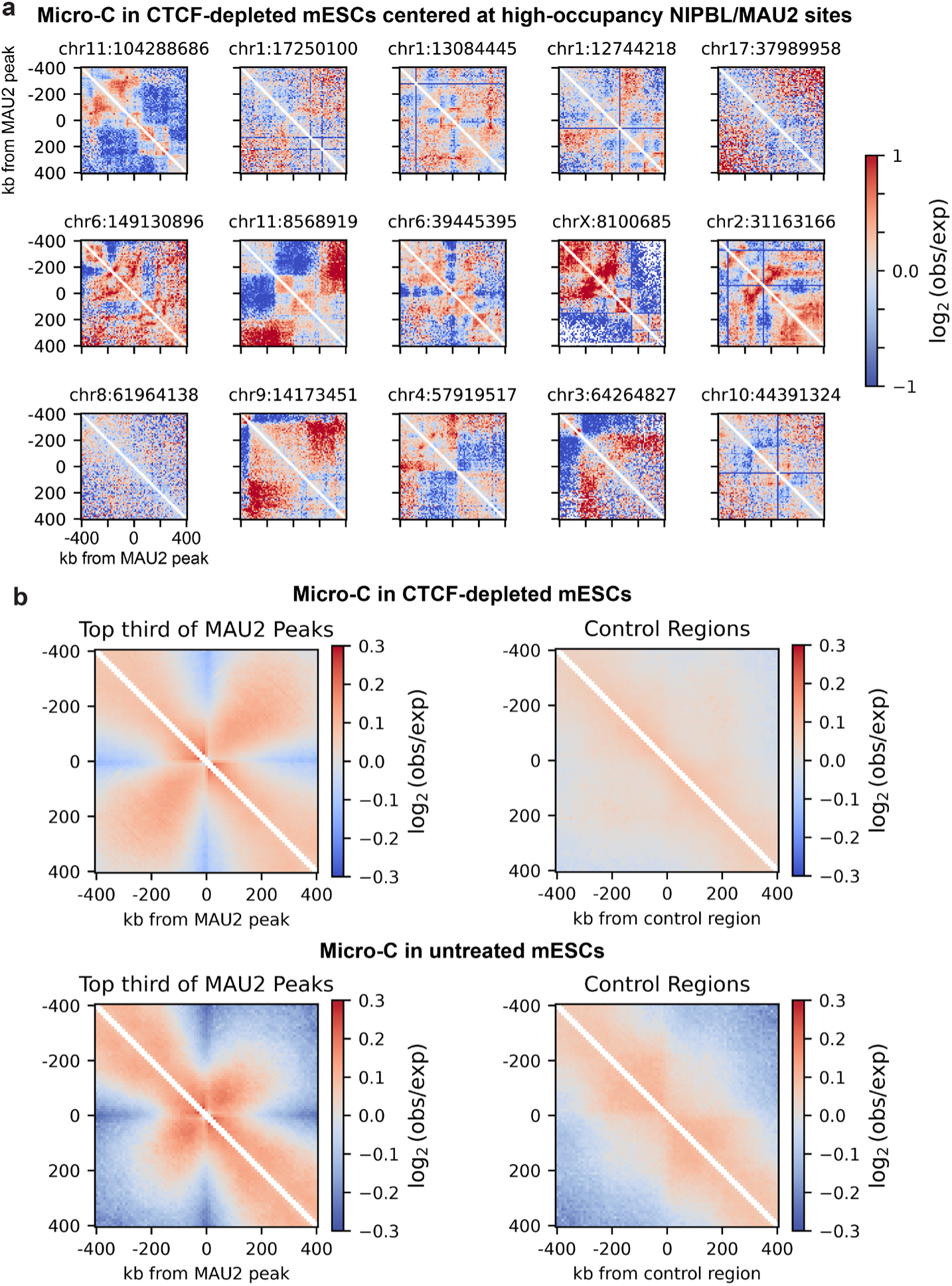
NIPBL/MAU2 sites only mildly change local chromatin folding. **a**, At 15 of the highest occupancy individual MAU2 sites, micro-C in CTCF-depleted cells does not show fountain patterns. Heatmaps are plotted at 10 kb resolution. **b**, A pile-up of micro-C data in CTCF-depleted or untreated ESCs^118^ at the highest-occupancy third of MAU2 sites shows a weak fountain pattern, while randomly chosen sites within the same TADs as the MAU2 sites do not.

**Extended Data Figure 9:**
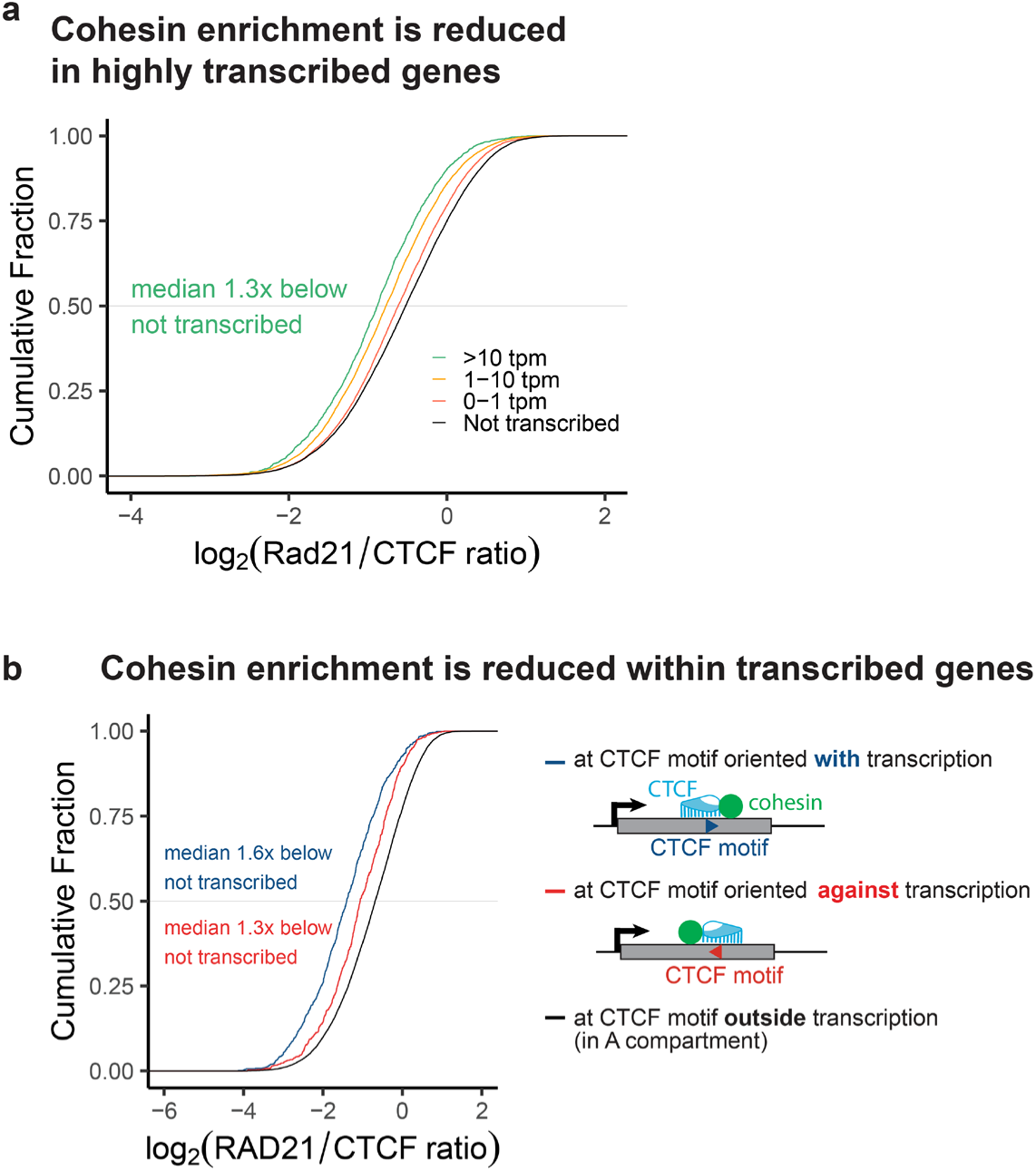
Cohesin traffic is impeded in actively transcribed loci. **a**, Cumulative plot shows lower RAD21/CTCF ratios for CTCF sites within gene bodies based on their transcription level. **b**, Cohesin enrichment is reduced at CTCF sites within GENCODE protein-coding genes with RNA-seq transcription >10 tpm^115^.

**Extended Data Figure 10:**
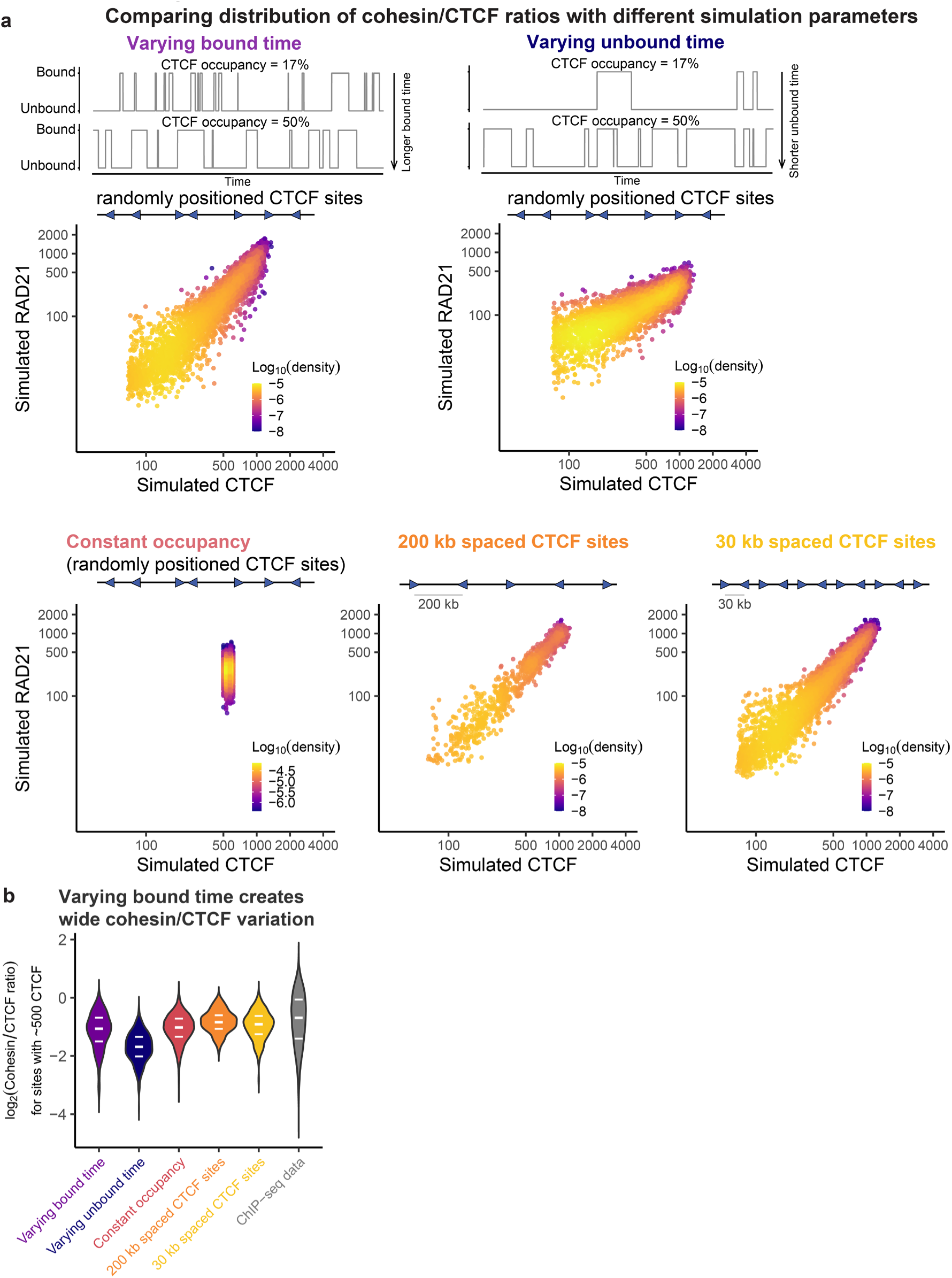
Comparison of CTCF/cohesin ratios using simulations with different parameters. **a**, Cohesin and CTCF occupancies at CTCF sites from simulations using different parameters for CTCF site occupancy and spacing. **b**, Distributions of CTCF/cohesin ratios for each parameter compared to ChIP-seq data.

**Extended Data Figure 11:**
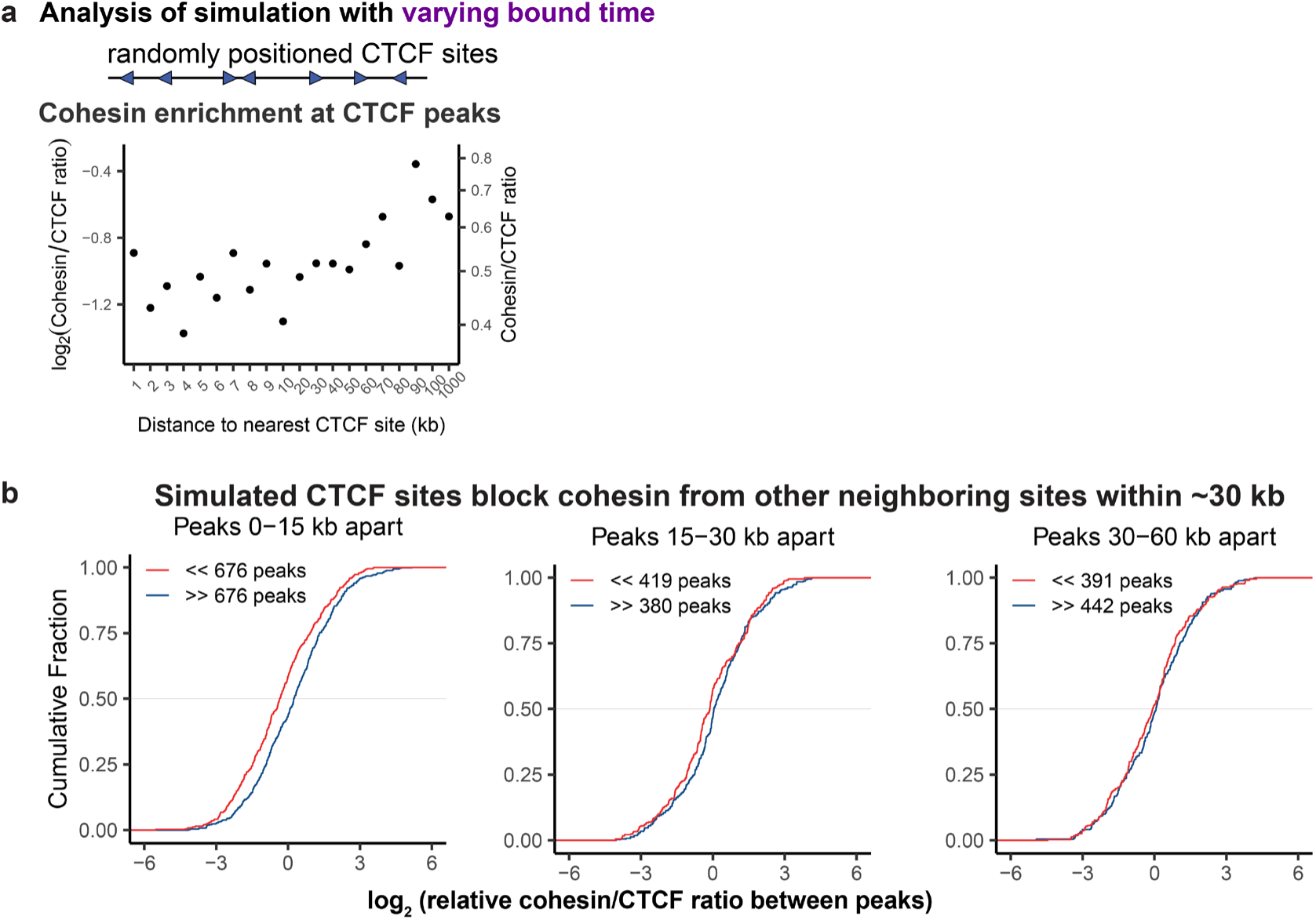
In simulations, neighboring CTCF sites block cohesin traffic. **a**, Simulation quantification showing that CTCF sites near other sites have lower cohesin occupancy (compare to ChIP-seq data in Fig. 6A, Extended Data Fig. 7D). **b**, Simulation quantification showing that CTCF sites block cohesin from other CTCF sites within ∼30 kb (compare to ChIP-seq data in Fig. 6C, Extended Data Fig. 7E).

**Extended Data Figure 12:**
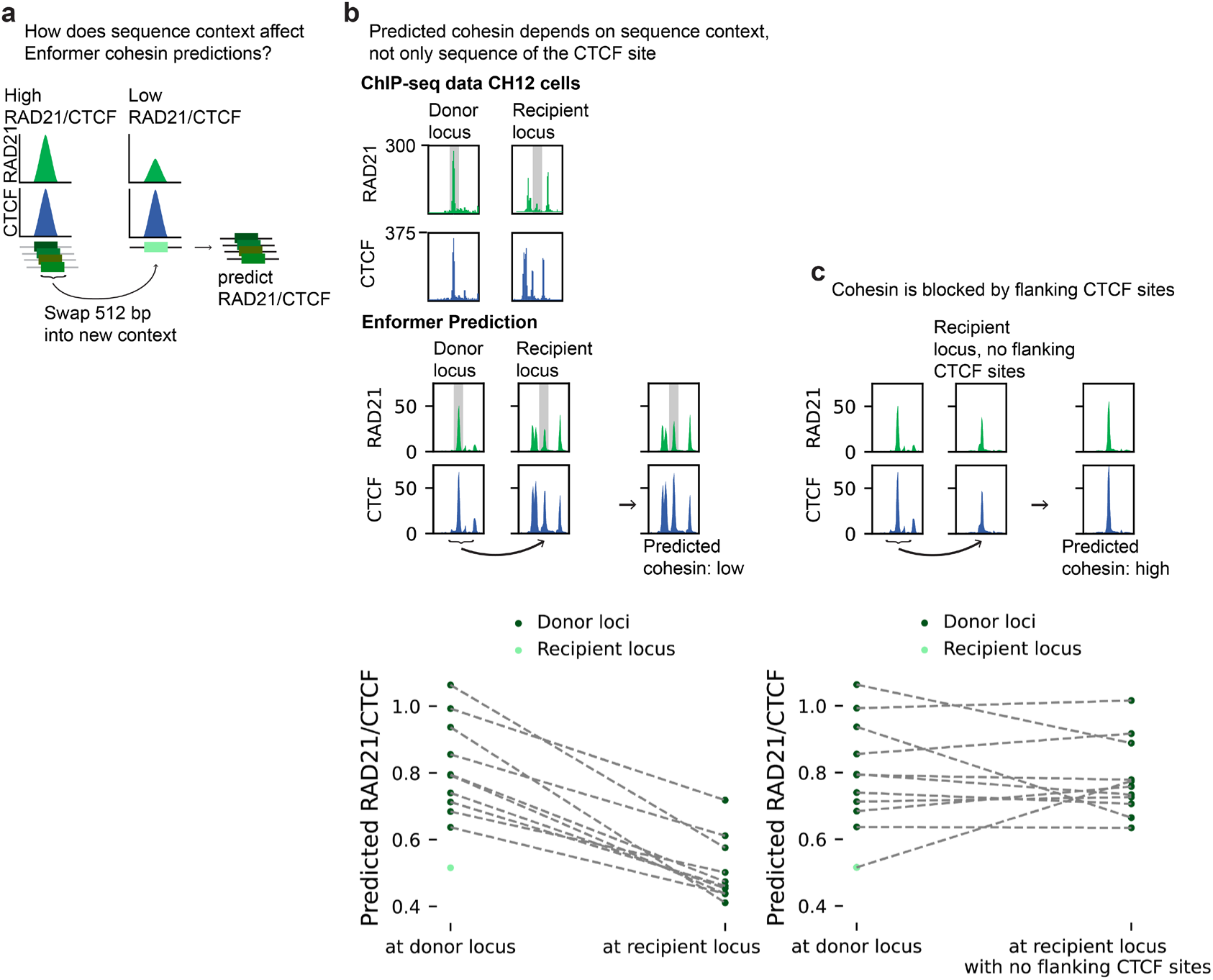
Enformer predicts that clustered CTCF sites alter cohesin accumulation. **a**, Using Enformer^120^, we selected sites with high predicted CTCF and RAD21 levels in CH12 cells (a mouse B cell lymphoma line), swapped 512 bp from the CTCF binding site into a recipient sequence context where a high-occupancy CTCF site has low RAD21 binding, and predicted binding for the chimeric sequences. **b**, The chimeric sequences show low predicted RAD21, indicating that the sequence context affects the predicted cohesin binding. Cohesin and CTCF binding ChIP-seq data from CH12 cells^115^ at the donor and recipient loci match the Enformer predictions. **c**, Without the CTCF motifs flanking the central site of the recipient locus, the predicted cohesin increased, indicating that Enformer predictions are influenced by CTCF motifs in the surrounding sequence.

**Extended Data Figure 13:**
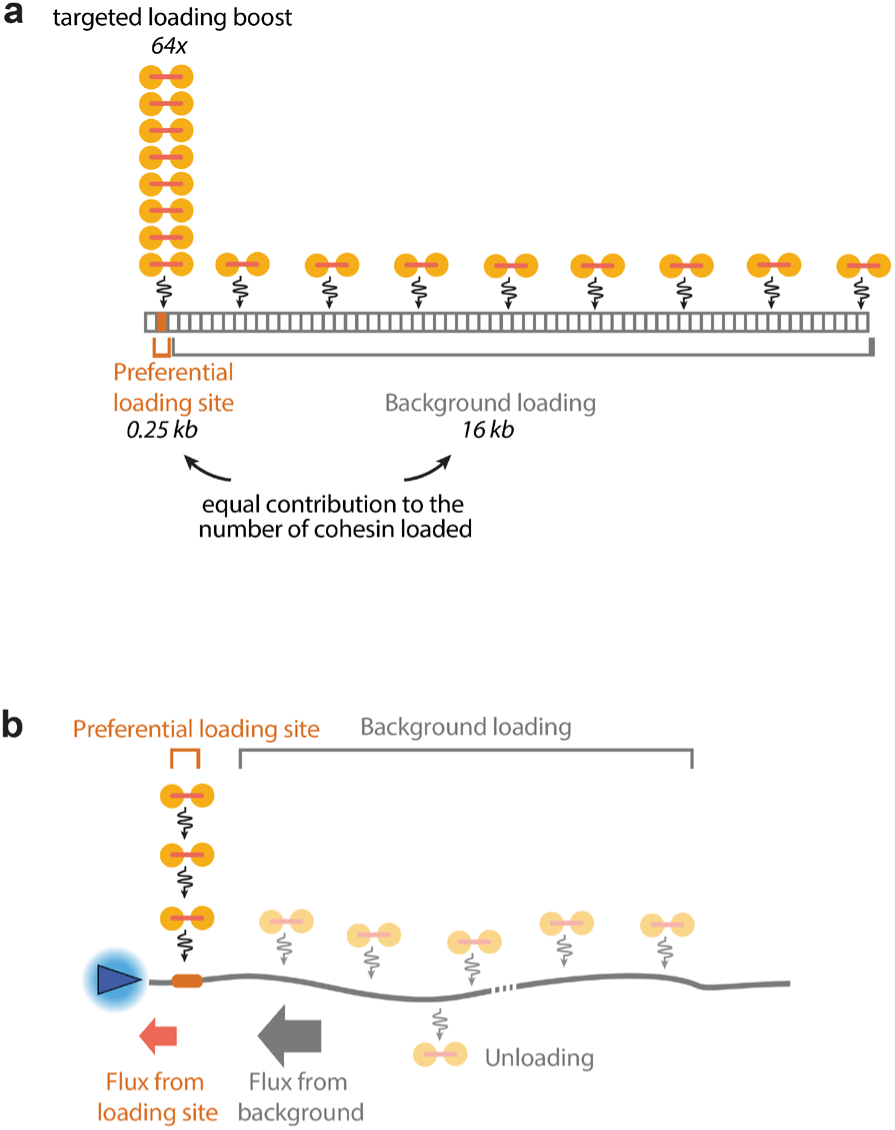
Cartoons comparing the contributions of targeted versus background cohesin loading. **a**, As an example, with a 64-fold targeted loading boost, a 0.25 kb loading site contributes to loading as much cohesin as a 16 kb background region. **b**, Cartoon illustrating the relative contributions of targeted and background cohesin loading to the flux of cohesin reaching a CTCF site. Even when a preferential loading site exists, its contribution is easily outweighed by cohesin loaded across the surrounding chromatin at the background rate.

