## Supplementary figures and images for "The role of cohesin loading at enhancers in the flux of loop extrusion and long-range transcriptional control"

### targeted loading extrusion movie

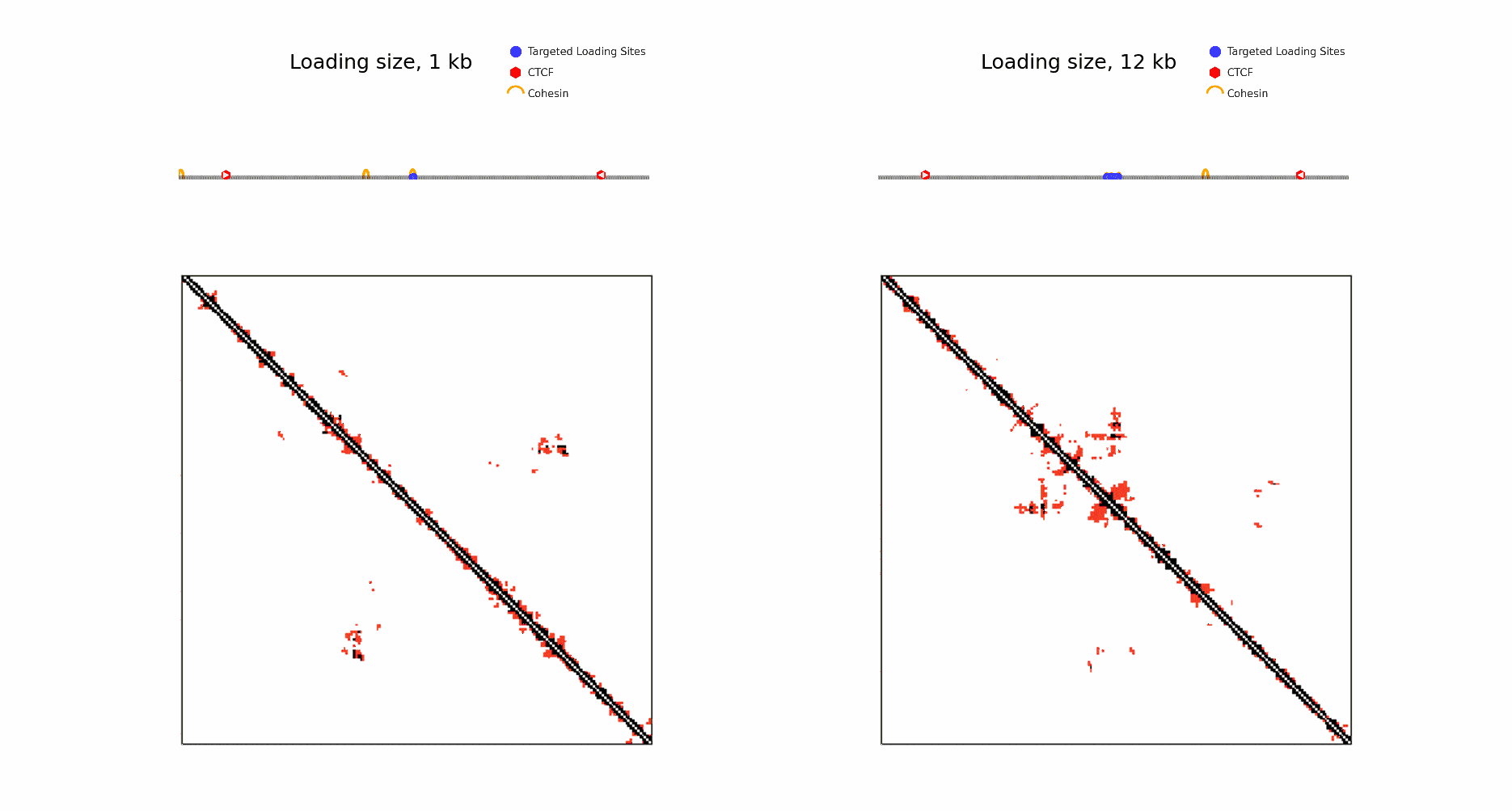
